# Hyperspectral open source imaging system

**DOI:** 10.1101/2024.06.21.600046

**Authors:** Jolyon Troscianko

**Affiliations:** Centre for Ecology & Evolution, University of Exeter

**Keywords:** Hyperspectral imaging, spectral imaging, light environment, artificial light at night, spectro-radiometry, ecology, environmental optical physics

## Abstract

The spatial and spectral properties of the light environment underpin the appearance of scenes and objects. As such its measurement is crucial for understanding many aspects of animal behaviour, ecology and evolution, along with many more applications in fields ranging from optical physics, agriculture/plant sciences, human psychophysics, food science, architecture and materials sciences. The escalating threat of artificial light at night (ALAN) presents unique challenges for measuring the visual impact of light pollution, requiring measurement at low light levels across the human-visible and ultraviolet ranges, across all viewing angles, and often with high within-scene contrast.

Here I present a hyperspectral open-source imaging system (HOSI), an innovative and low-cost solution for collecting full-field hyperspectral data. The system uses a Hamamatsu C12880MA micro spectrometer to take single-point measurements, together with a motorised gimbal for spatial control. The hardware uses off-the-shelf components and 3D printed parts, costing around £350 in total. The system can run directly from a computer or smartphone with a graphical user interface, making it highly portable and user-friendly.

The HOSI system can take panoramic hyperspectral images that meet the difficult requirements of ALAN research, sensitive to low light around 0.001 cd.m^-2^, across 320-880nm range with spectral resolution of ∼9nm (FWHM) and spatial resolution of ∼0.2 degrees. The independent exposure of each pixel also allows for an extremely wide dynamic range that can encompass typical natural and artificially illuminated scenes, with sample night-time scans achieving peak-to-peak dynamic ranges of >50,000:1.

This system’s adaptability, cost-effectiveness, and open-source nature position it as a valuable tool for researchers investigating the complex relationships between light, environment, behaviour, ecology, and biodiversity, with further potential uses in many other fields.

## Introduction

Light carries a wealth of information about the world around us; primarily conveyed in its intensity, spectral composition and angular direction. Taken together, this spatio-spectral information forms the basis of animal vision; guides behaviour, and ultimately impacts animal evolution via natural or sexual selection (Endler, 1993). Spatio-spectral information is also used by a range of sensing/measurement systems, including optical physics (Akkaynak et al., 2017), plant ecology and agriculture (Imran et al., 2023; Youngentob et al., 2012), food technology, geology/mineral science, and astronomy (Hänel et al., 2018). Quantifying the light field is also critical for accurate modelling of object and scene appearance, linking fields as diverse as human psychophysics, visual ergonomics, lighting design and architecture, and digital creative industries (Morimoto et al., 2019; Yu et al., 2023).

The light environment is fundamental to all aspects of visual ecology; from behaviour to evolution, and can even drive speciation (Price, 2017; Seehausen et al., 2008). Natural scenes vary substantially in their spatio-spectral properties (Párraga et al., 1998), largely driven by the type of illuminant (such as sunlight, moonlight, starlight, bioluminescence or artificial light), and also how the light propagates through the environment as it is absorbed, scattered and reflected. Factors such as scattering (e.g. Rayleigh scattering or oceanographic backscattering), cloud/fog/forest canopy/algal absorption, and Snell’s window determine the intensity, spectral composition and spatial structure of the light environment (Akkaynak & Treibitz, 2018; Bohren, 1987; Cronin et al., 2014; Tidau et al., 2021). For example, a cloudy/foggy day will result in isotropic illumination (spatially uniform illumination), reducing shadows and contrast. Direct light (such as sunshine) will result in interactions between the habitat’s 3D structure and lighting angle to create strong shadows likely to increase the scene’s visual complexity and make visual search more difficult (Xiao & Cuthill, 2016). The light’s directional properties will also affect critical anti-predator defences such as countershading, which is dependent on the intensity of an animal’s self-shadows (Cuthill et al., 2016).

Artificial light at night (ALAN) is increasingly recognised as a global threat to biodiversity (Rich & Longcore, 2013; Sanders et al., 2021), and over the past 18 years it’s intensity has more than quadrupled (Kyba et al., 2023). ALAN can modify animal and plant behaviour and phenology through a number of mechanisms that vary in their spectral sensitivity ranges (Briolat et al., 2021; Davies et al., 2013; Sanders et al., 2021). Existing data on the global extent of ALAN are largely limited to satellite-based data that measure upwelling light on cloud-free nights, so are unable to account for atmospheric effects and horizontally propagated light. Marine systems are also increasingly recognised as vulnerable to the effects of ALAN, and measurements in these conditions are even more difficult (Tidau et al., 2021). Many existing studies are also unable to measure the spectra most relevant to animal and plant responses, or weigh the likely impacts to non-human receivers (Kyba et al., 2023; Sánchez de Miguel et al., 2022). Measuring skyglow – which is particularly strong under cloud-cover – is increasingly recognised as a major factor in the intensity of ALAN (Kyba et al., 2023; Stöckl & Foster, 2022), that also causes substantial shifts in the light’s directional structure. Recent work has also highlighted the potential for the directional structure of lighting to impact animal anti-predator defences, with diffuse light causing a mismatch between colour change and background selection in a marine isopod (Bullough et al., 2023). Quantifying the light environment’s spatial and spectral properties is therefore of particular importance for ALAN research.

A number of existing methods have been developed for quantifying spatio-spectral information. No clear consensus exists on the distinction between multispectral and hyperspectral imaging systems (Polder & Gowen, 2021), however, I propose that ‘hyperspectral’ should be reserved for systems that operate at substantially higher spectral resolution than the full-width at half-maximum (FWHM) of typical opsin responses. Multispectral techniques such as wide-angle photography can be used to photograph whole scenes, with calibration allowing for conversion to absolute radiance (Nilsson & Smolka, 2021). Further recent work has used similar methods to model non-human low-light visual dynamics to ALAN and sky glow in particular (Stöckl & Foster, 2022). This method enables rapid data collection using affordable equipment, with high spatial resolution and moderately high dynamic range. While exposure bracketing can increase the effective dynamic range of such systems, in practice the optical train’s internal reflections and modulation transfer function will often result in under-estimates of true scene dynamic ranges. Photography-based methods also have limited spectral resolution (human RGB only, and will not readily work for ultraviolet ranges due to the limitations of wide-angles lenses). Multispectral and hyperspectral camera systems that use pass filters are a good compromise between spatial and spectral resolution under natural lighting for many tasks (Párraga et al., 1998; Troscianko & Stevens, 2015). However, as above, wide-angle lenses cannot operate over the ultraviolet-visible range, and the spectral resolution is not sufficient for ALAN work that often involves complex emission spectra [resulting in metamerism in low-spectral resolution systems (Troscianko, 2022)]. These camera systems also require moderately complex calibration to convert pixel values to absolute radiance. Hyperspectral cameras can be used to image reflective spheres in order to build up whole-scene radiance with higher spectral resolution than that offered by camera-based systems (Morimoto et al., 2019). This technique can provide moderately high spectral and spatial resolution, however there are a number of drawbacks. Notably the dynamic range of existing hyperspectral cameras is comparatively low (because the optics and exposure control must now operate over a wider band of spectra), and they struggle to measure the vast dynamic range typical of natural scenes. Absolute sensitivity is also typically low, making them poorly suited to low-light situations (such as ALAN research). Capture and calibration is also complex as it requires taking and stitching together multiple scans from different angles to build up a whole scene on top of the above calibrations required to convert to absolute radiance. Hyperspectral cameras are also comparatively expensive (typically >£20k for human-visible range, or considerable more for UV-visible systems), and far beyond the resources of many researchers. Motorised (gimbal-mounted) spectroradiometers or imaging telescopes are a final technique, such as the ASTMON systems for quantifying night-time lighting for astronomy (Hänel et al., 2018). Nevertheless, such systems are not typically affordable, accessible, or easily adapted to measuring high dynamic range light environments. I therefore developed a low-cost, high-sensitivity imaging system capable of producing spectral radiance or relative reflectance images of whole-scenes across wavelengths relevant to any visual system.

## Materials and Methods

All code, calibration data and sample scans are provided as supplementary information with this submission, however, see the Github project for the latest updates: https://github.com/troscianko/HOSI/

### Hardware

The HOSI system was developed from OSpRad; an open-source spectroradiometer designed for low-light spectral radiance and irradiance measurements (Troscianko, 2022). Both systems are built around the Hamamatsu C12880MA micro spectrometer, which has comparatively high sensitivity (down to 0.001 cd.m^-2^), a high spectral resolution with 288 photosites (typical FWHM of 9nm, and maximum FWHM of 15nm), and a spectral range of ∼320-850nm that encompasses effectively all animal phototransduction-based visual systems, and the PAR/near-IR ranges required for plant-based measurements. The component is low-cost (approximately £220 from Hamamatsu UK), and is typically used in medical equipment, but can be interfaced through a customised open-source microcontroller. For further details on how the microcontroller interfaces with the C12880MA, see Troscianko (2022), and the HOSI’s C++ Arduino code. Parts were printed in matt black PLA (Polylactic acid, SUNLU Matte PLA 1.75mm Printer Filament, Amazon UK, ∼£18, figure 1c). It is essential the the printed plastic does not transmit near-infrared light (e.g. avoid black PET, Polyethylene terephthalate). See figure 1 for examples.

**Figure 1.**
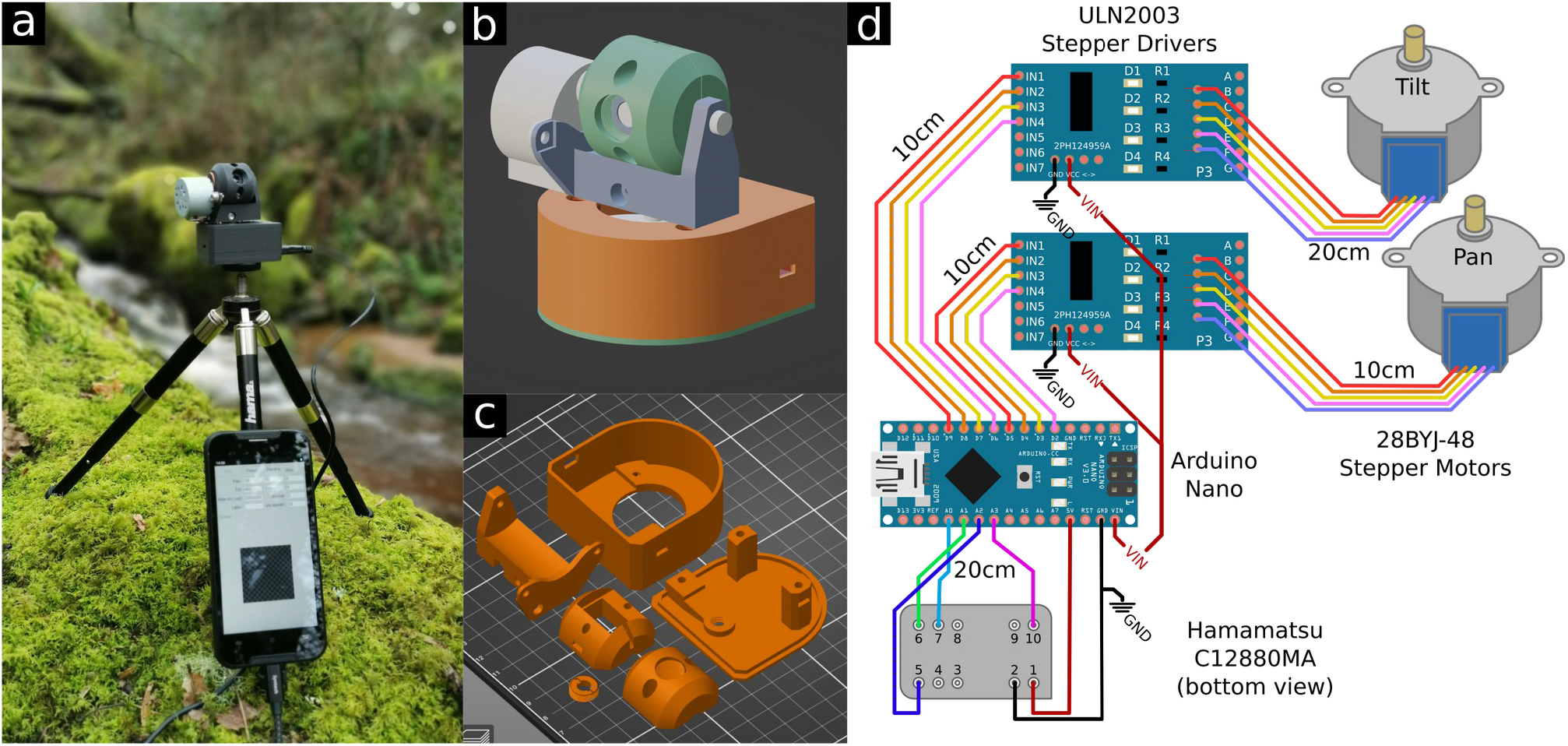
3D parts and wiring. a) Photograph of a HOSI unit in the field, running from a smartphone. b): 3D assembly; c): parts arranged for 3D printing; d): wiring diagram and required wire lengths. GND (ground) is the shared negative voltage, VIN is the 5v output from the Arduino that is delivered via the USB cable. Note that the spectrometer should be connected to the Arduino’s regulated 5v output (as shown), not VIN that has a slightly lower voltage due to a protective diode in the Arduino.

Focusing is achieved using a biconvex fused silica lens with 10mm focal length, and 6mm diameter (LB4280; Thorlabs Ltd. ∼£75). This is an off-the-shelf lens that provides sufficient spatial detail for most imaging and light environment measurement tasks. The lens is placed into a 3D-printed threaded holder that can be screwed into the HOSI housing for manual focusing from infinity to roughly 30mm, providing a roughly 0.5 degree acceptance angle. A 3D printed tool allows for easy focusing, and a user can make marks on this tool to denote the positions of different focal depths. This system is well suited to low-light applications, but will saturate under strong daylight light intensities, so light reducer aperture 3D printed parts are also provided for daytime use (note that the spectral sensitivity measurements below must be repeated if a light reducer is used).

Hyperspectral images are acquired by using a 3D printed gimbal that pans and tilts the spectroradiometer with stepper motors. 28BYJ-48 stepper motors were selected for their high angular resolution of 2048 steps per revolution, an ability to run from a USB (5v) source, their availability, and low cost. The spectrometer and stepper motors are controlled by an Arduino Nano microcontroller; this in turn interfaces through serial connection with a computer or smart phone. The housing uses 3D printed parts and can be assembled with basic soldering skills (figure 1d).

### Calibration

#### Linearisation

Calibration broadly followed Troscianko (2022). While the uncalibrated system has a good linear response to radiance across most of its range (R^2^ values typically ∼0.996-0.998 using the below linearisation protocol), low count values (below roughly 1% of maximum) over-estimate radiance relative to the middle and upper ranges. To make linearisation more straightforward I developed a Python script (file: Calculate_HOSI_linearisation_curve_0.6.py) that automates the process by taking 22 measurements with exposures ranging from 0.005 to 1.5 times the auto-exposed integration time. To use the script, the HOSI should be placed in front of a surface that is illuminated with a source that will remain stable in spectral output and intensity for roughly one minute. For most reliable results this should be performed in a dark room to eliminate unwanted light ingress during the dark measurement. Almost any stable non-flickering source can be used in principle, such as conventional non-flickering LED room lighting, or a display screen. Ideally the intensity should use upper integration times of around 200,000-800,000 microseconds to maximise the dynamic range. The script removes any saturated (over-exposed) values, and selects the spectral peak of the linearisation spectrum. Values at this peak wavelength are then compared between known exposure times, comparing observed and expected (linear) responses. A Gaussian smoothing function (σ=3) is applied to the spectral recordings to reduce noise, and dark count values are subtracted from light counts. The script then uses least squares regression to fit the results to a linearisation equation, and outputs the model coefficients together with R^2^ values and log-log axis plot. This linearisation modelling typically increases R^2^ values to >0.999, with performance at low-values notably improved.

#### Spectral sensitivity

Stable illumination in the UV-visible range was provided by an Osram XBO R 100 W/45 bulb (∼$435), driven by a USHIO SMARTARC™ HBX-76 igniter and ballast (∼$500), both sourced from LabGear Inc. The lamp was fitted to a stainless steel hemispherical reflector (made from an IKEA BLANDA bowl £5). The XBO lamp provides a highly focused beam of light that is not suitable for diffuse illumination. To diffuse the light source a 80mm diameter disk of 1mm thick virgin PTFE sheet was fitted roughly 100mm from the bulb. A second, larger diffuser disk 300mm diameter made from 0.5mm thick virgin PTFE was then used for further diffusion mounted roughly 150mm from the bulb. A centrifugal fan at the base of the lamp provides cooling to the bulb and diffusers (the ballast provides a 12v output for this purpose). The resulting lamp provides unfocused, roughly cosine-shaped throw and is far better suited to typical calibrated photography/hyperspectral imaging than most stable xenon sources available from commercial suppliers, while being considerably cheaper. However, care must be taken to house and use the system in a safe way given its ionising radiation emissions, high voltage spikes at ignition, and potential for explosion of the high-pressure XBO lamp. Spectral sensitivity calibration was achieved by taking a hyperspectral image of a Spectralon 99% reflectance illuminated by the stable Xenon source (above), and comparing this to a calibrated spectral radiance measurement. The illuminant was directly above the standard (90 degrees from the surface), while measurements were taken from approximately 45 degrees using the HOSI system, and a NIST-tracable Jeti Specbos UV-1211 spectroradiometer. Raw count data were then extracted from the hyperspectral image file (averaged across the 35 pixels that recorded the white standard), together with count data of the dark measurements taken before and after the white standard, matching the same integration. Average dark counts are then subtracted from average light count data at each wavelength. A Python script (file: Calculate_HOSI_spectral_sensitivity_0.2.py) can then be used to calculate the spectral sensitivities from these count data and calibrated spectrum, together with the integration time (in microseconds) and the linearisation coefficients (measured above).

Calibration data must be saved to the calibration_data.txt file in the same directory as the GUI’s Python code. This file specifies the photosite wavelength calculation coefficients (provided with each Hamamatsu C12880MA chip by the manufacturer), linearisation model coefficients, and spectral sensitivity functions for each unit. Each HOSI unit can have its own unit number that is specified when uploading the firmware, and the calibration_data.txt file allows for multiple units to be added (so labs can use multiple units across smartphones/computers without risk of using the wrong calibration data). All testing was performed at standard room temperatures of around 19-20 degrees. Sensor performance may vary at temperatures well outside this range, and users should calibrate and test their units under typical use scenarios.

### Firmware

The Arduino Nano microcontroller uses custom written code (‘firmware’, file: Arduino_HOSI_1.05.ino, written in Arduino C++), that was developed from the OSpRad project as above (Troscianko, 2022), however it has a number of major changes to optimise the system for speed while maintaining high-dynamic range, and providing gimbal positional control. This firmware is uploaded to the Nano using Arduino IDE software.

#### Dark measurement

The spectrometer uses a CMOS sensor, which converts absorbed light at photocells into a potential difference (higher voltage for more light received), an analogue-to-digital-converter on the Arduino Nano then converts this to a digital count. However, each photocell also has a “dark current” associated with the black point, and this varies with factors such as supply voltage and sensor temperature, so must be measured regularly. The HOSI system automatically makes regular (and user customisable) dark measurements by rotating the spectrometer down into the gimbal housing at the end of a panning scan, to a position designed to block light. These dark measurements are stored in the raw output for subsequent calculation of absolute spectral radiance.

#### Exposure control

Exposures are calculated independently at each gimbal position, with integration times starting from the minimum of 500 microseconds, and increasing in octaves until an exposure is saturated in any wavelength, or the user-specified maximum integration time is reached (default of 2 seconds, maximum 30 seconds). The firmware then sends the previous (non-saturated) exposure to the app (below), and proceeds to the next gimbal position. The theoretical dynamic range of the system within a single hyperspectral image is therefore approximately 1:60,000 based on integration times alone, and considerably higher when considering the dynamic range of the sensor itself.

#### Scanning behaviour

The user can specify the start point, stop point and resolution (steps per measurement) for the pan and tilt axes independently. Panning allows measurements across 360 degrees (azimuth), and tilt from approximately -20 degrees 90 degrees (elevation). The stepper motor gears introduce a small amount of play that the firmware overcomes by overshooting and returning to position when starting a new panning scan.

#### Projection

HOSI uses a Mercator projection to link each pixel’s azimuth and elevation to 2D image x and y coordinates respectively. This would result in higher resolution measurements near the poles (high elevation measurements), and as such the firmware progressively skips redundant pixel scans near the poles, interpolating measurements for efficient whole-scene scans.

#### Spectral Compression

Spectral data can be compressed by selecting an averaging value that combines neighbouring photosite count data over a user-specified window size. For example, a value of 2 halves the number of output data without reducing the effective spectral resolution (given the C12880MA’s ∼9nm FWHM resolution). Window sizes greater than 2 will result in loss of spectral resolution.

### Graphical User Interface (GUI) Application

The HOSI unit receives both power and communication via its USB connection with a computer or smartphone, making the system highly portable and flexible. The graphical user interface application (app, file: HOSI_GUI_0.1.45.py) was written in Python 3.10.6, and provides straightforward access to control a HOSI unit and save the output data to the computer/smartphone (see figure 2). The code can run on computers with Python, or on Android smart phones using the free Pydroid app. The app uses the calibration data (above) to calculate radiance for each pixel (*Le* [W.sr^-1^.m^-2^.nm^-1^]), which is saved in the output file along with raw data, an sRGB image of the scan, a user-specified label, and time-stamp.

**Figure 2.**
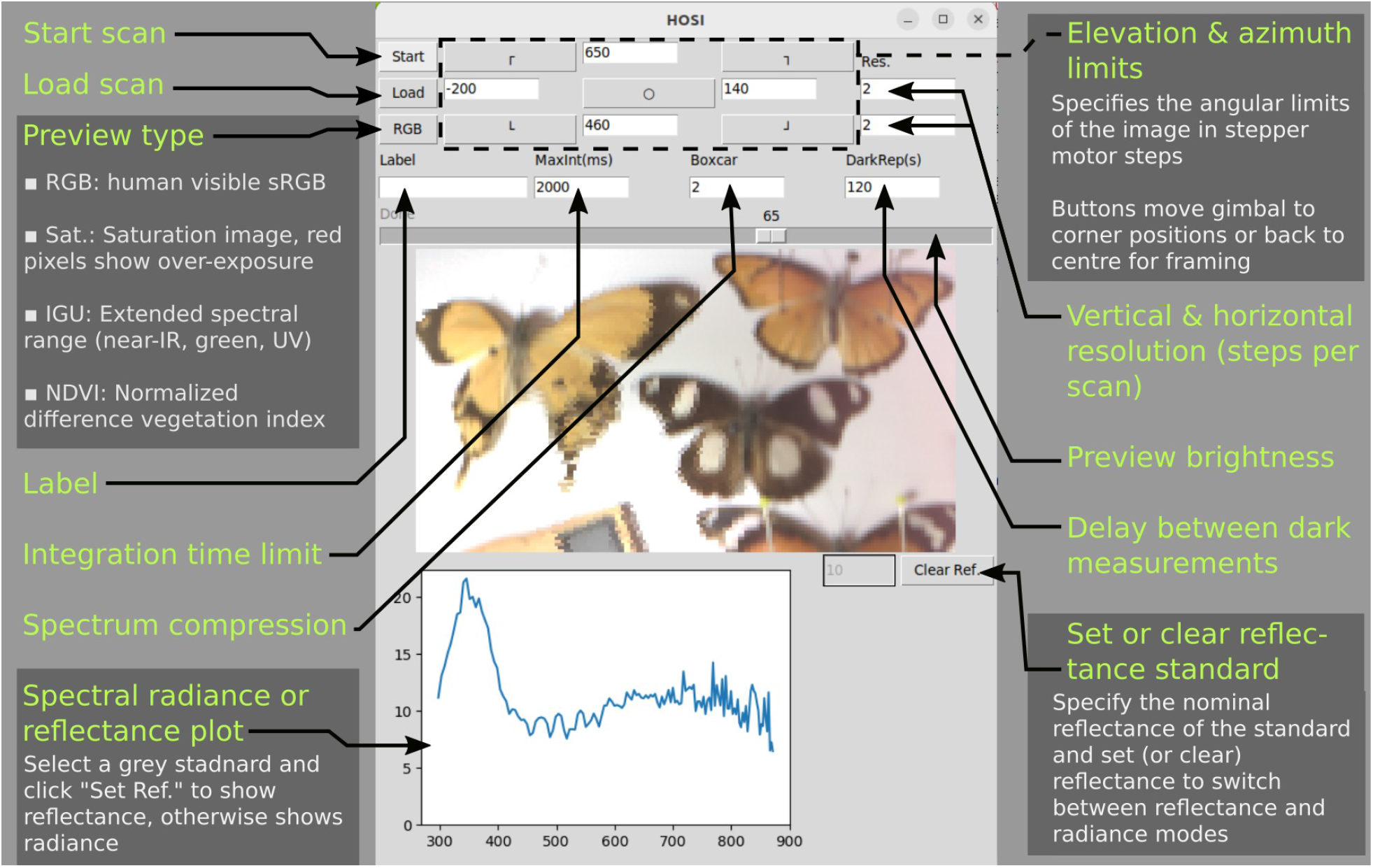
Graphical user interface (GUI) layout, showing the available functions and a scan of butterflies. The spectrum shows a selection from a patch of wing that appears black to human vision, but has a UV peak visible to many other animals. This scene was illuminated by an Exo-Terra Sunray lamp.

The app supports multiple preview formats, including: RGB (conventional sRGB image), an extended spectral range image that uses near infrared, green (CIE Y), and ultraviolet in place of RGB values respectively, a saturation preview mode that highlights any pixels where saturation has occurred through over-exposure, and an NDVI image for vegetation.

Each pixel’s spectral radiance (*Le*) can be plotted by tapping/clicking on it in the preview. Output can also be viewed as relative reflectance by selecting a pixel containing a grey or white standard and clicking “Set Ref.”. Once set, pixel spectrum plots now show relative reflectance (% relative to the standard), and the preview image has the white-balance set.

## Discussion

The hyperspectral open-source imaging system (HOSI) presented in this study addresses the growing need for affordable and versatile tools for quantifying spatio-spectral properties of light environments. In particular it offers high spectral and spatial resolution, high sensitivity, and a far larger dynamic range than existing ultraviolet to near infrared systems (because each pixel has an optimal exposure and glare-shielded optical train). Figure 3 shows a night-time scene measured with the HOSI system, demonstrating how individual bulb technologies can be identified, while also highlighting the large dynamic range present (with the highest spectral peak over 50,000 times higher than the lowest spectral peak in the scene). Moreover, the system is small and portable, capable of being operated with a smart phone, and has comparatively straightforward calibration compared to existing solutions, with output data that are largely pre-processed.

**Figure 3.**
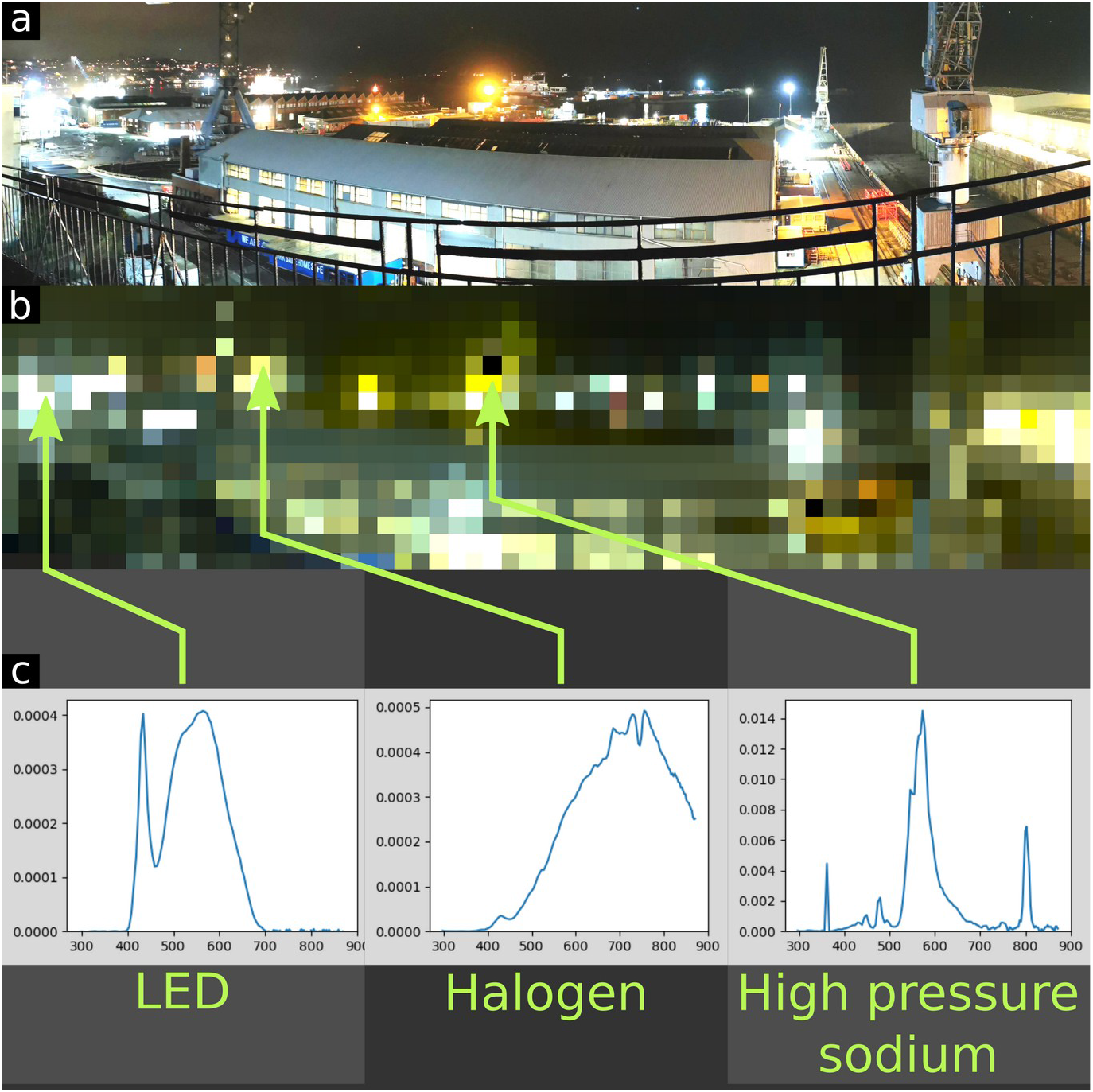
Example hyperspectral scan of Falmouth (UK) docks at night. a): panoramic photograph of the scene; b): HOSI’s hyperspectral RGB preview image; c): radiance plots created by the HOSI GUI at chosen pixels (x-axis: wavelength [nm]; y-axis: linear radiance *Le* [W.sr^-1^.m^-2^.nm^-1^]). These spectra illustrate how individual light sources can be identified and compared. The highest spectral peak in the brightest pixel of this hyperspectral image is approximately 50,000 times higher than the peak in the darkest pixel. The full dynamic range (highest peak to lowest trough) is considerably greater, and can be maximised further with longer exposure times to reduce noise.

While the HOSI system offers significant advantages, a notable limitation is the time it takes to scan high-resolution images, particularly under low-light. As such the system is not suited to instantaneous imaging of scenes with fast-changing illumination or moving objects. However, high spatial resolution is typically not required for quantifying the light environment, and in many cases movement or spatial changes may be statistically averaged out in large datasets. Moreover, the system’s flexibility allows researchers to adjust elevation and azimuth resolutions independently based on their specific needs, balancing time constraints with spatial requirements. For example, Nilsson & Smolka (2021) use whole-scene photographic techniques with high spatial resolution, but the data are subsequently converted to vertical transects that a HOSI system could readily replicate through transects. When used as a hyperspectral imaging camera, the HOSI system can be used in the lab with a stable full-spectrum light source for generating high spectral resolution images where capture time is not limiting.

The HOSI’s adaptability, and open-source nature contribute to its potential for environmental sensing and light environment quantification in a wide range of scientific disciplines. Furthermore, its affordability allows the system to be used in high-risk environments (aerial or underwater sensing (Tidau et al., 2021)), or at increased scope by deploying multiple systems. The project also paves the way for future innovations in spatio-spectral data measurement systems that require co-ordination of spectrometry with actuators, such as automated goniometry for quantifying iridescence (Gruson et al., 2018), or automated sensing systems in medicine or food testing.

## Supporting information

HOSI supplementary files including code and 3D printer models

## Data Availability

All code, 3D models, calibration data, and sample scans are available at the project’s GitHub page: https://github.com/troscianko/HOSI/, released under a GNU General Public License v3.0.

