## Supplementary material for "Hyperspectral open source imaging system": HOSI supplementary files including code and 3D printer models: Arduino_HOSI_1.05.ino

```
/*

_________________________HOSI Arduino Firmware_____________________________

 Written by: Jolyon Troscianko
 initial release: 19/06/2024
 License: GNU General Public License v3.0
 
 
 Manual control:
 The HOSI can be controlled automatically by the HOSI_GUI Python software. however, the functions can
 also be accessed manually via a serial port (check the baud, default 115200)
 
 tilt: send "l" followed by a number e.g. "l100" or "l500" note that zero is straight down, do not use negative values
 pan: send "p" followed by a number e.g. "p100" or "p-1000" note that zero is straight ahead
 manual integration time: send "t" followed by microsecond number
 radiance measurement: send "r" and it will take a radiance measurement
 hyperspectral measurement: send "h" followed by comma-delineated values as follows:
 panLeft,panRight,panResolution,tiltBottom,tiltTop,tiltResolution,maxIntegrationTime(microseconds),boxcar,darkRepeatTimer(milliseconds)
 e.g.: "h-200,200,10,400,600,10,2000000,2,120000"
  
 
 Modify the unit number below if desired, and upload this script to an Arduino Nano or similar.
 
 
 */


#include <AccelStepper.h>


int unitNumber = 9; // Change this number to whatever you want, and link this to the calibration data in calibration_data.txt


#define TRGpin A0
#define STpin A1
#define CLKpin A2
#define VIDEOpin A3

#define nSites 288 // 
uint16_t data[nSites] [2];
int dataLoc = 0;

int delayTime = 1;
long intTime = 100;
long prevIntTime = 100;
long maxIntTime = 3000000; // maximum integration time for auto & manual measurement milliseconds
int minIntTime = 50; //the sensor doesn't work below a certain numebr of scans... it needs three cycles with the ST pin high, even then you need to add about 378 microseconds to the exposure
int intStep = 400;
long manIntTime = 0;
int satN = 0; // number of bands over-exposed
int prevSatN = 0;
int satVal = 998; // over-exposure value

// used to control the reduction of redundant measurements at high elevations
int panoSteps[] = {913, 959, 988, 1005};
int panoSpaces[] = {2, 4, 8, 16};

long hyperVals[9];
long panVal = 0;
long tiltVal = 0;
int darkLight = 1; // 0= dark measurement, 1=light measurement
long darkRepeat = 30000;// repeat dark measurement every few seconds/minutes

AccelStepper stepper_pan(4, 3, 5, 2, 4); // 8=HALF4WIRE, then input pin 1, 2, 3, 4
AccelStepper stepper_tilt(4, 7, 9, 6, 8); // 8=HALF4WIRE, then input pin 1, 2, 3, 4

int boxcar = 1;


void setup() {
  stepper_pan.setMaxSpeed(500.0); // 800 for half-step
  stepper_pan.setAcceleration(5000.0);
  stepper_pan.disableOutputs();

  stepper_tilt.setMaxSpeed(500.0);
  stepper_tilt.setAcceleration(5000.0);  
  stepper_tilt.disableOutputs();
      
  //Set desired pins to OUTPUT
  pinMode(CLKpin, OUTPUT);
  pinMode(STpin, OUTPUT);

  digitalWrite(CLKpin, HIGH);
  digitalWrite(STpin, LOW);

  Serial.begin(115200); // Baud Rate set to 115200, 500000 seems to work well for nano with PC, 230400 and 256000 doesn't work for android. interestingly it still works ok with 57600
  while (! Serial); // Wait untilSerial is ready
  readSpectrometer();
  resetData();
  
}


void readSpectrometer(){

  // Start clock cycle and set start pulse to signal start
  digitalWrite(CLKpin, LOW);
  delayMicroseconds(delayTime);
  digitalWrite(CLKpin, HIGH);
  delayMicroseconds(delayTime);
  digitalWrite(CLKpin, LOW);
  digitalWrite(STpin, HIGH);
  delayMicroseconds(delayTime);

  // microseconds
     unsigned long cTime = micros(); // start time
     unsigned long eTime = cTime + intTime; // end time
  
  //Sample for a period of time
 // for(int i = 0; i < 15; i++){ //orig 15
     while(cTime < eTime){
         digitalWrite(CLKpin, HIGH);
         delayMicroseconds(delayTime);
         digitalWrite(CLKpin, LOW);
         delayMicroseconds(delayTime);
         cTime=micros();
      } 

  //Set STpin to low
  digitalWrite(STpin, LOW);

  //Sample for a period of time
  for(int i = 0; i < 88; i++){ //87 aligns correctly

      digitalWrite(CLKpin, HIGH);
      delayMicroseconds(delayTime);
      digitalWrite(CLKpin, LOW);
      delayMicroseconds(delayTime); 
      
  }

  int specRead = 0;
  satN = 0;
  for(int i = 0; i < nSites; i++){

      specRead = analogRead(VIDEOpin);
      // second read shoudl stablise the multiplexer and give more accurate read
      //delayMicroseconds(delayTime);
     // specRead = analogRead(VIDEOpin);
      data[i][dataLoc] += specRead;
      if(specRead > satVal)
        satN ++;
      
      digitalWrite(CLKpin, HIGH);
      delayMicroseconds(delayTime);
      digitalWrite(CLKpin, LOW);
      delayMicroseconds(delayTime);
        
  } 
}


void resetData(){
  for (int i = 0; i < nSites; i++)
    data[i][dataLoc] = 0;
}

void switchDim(){
  if(dataLoc == 0)
    dataLoc = 1;
  else
    dataLoc = 0;
}

void radianceMeasure(){

  // reset all data
  dataLoc = 1;
  resetData();
  dataLoc = 0;
  resetData();

  // ----------------- AUTO EXPOSURE-----------
  if(manIntTime == 0){ 
    intTime = minIntTime; // microsecond exposure
    readSpectrometer(); // read to dim0
    //data[0][dataLoc] = -1; // debugging
    prevSatN = satN;
    prevIntTime = intTime;
    intTime = intStep;
    if(satN > 0)
      switchDim(); // required if first exposure is over-exposed

    while(satN == 0 && intTime < maxIntTime){
      switchDim();
      resetData();
      readSpectrometer();
      //data[0][dataLoc] = intTime;// debugging

      if(satN == 0){
        prevSatN = satN;
        prevIntTime = intTime;
        intTime = intTime*2;
      }

    } //while

    //if(prevIntTime > -1) // if first exposure isn't over-exposed
      switchDim(); 
      
  //------------- MANUAL EXPOSURE-------------
  } else { 
    intTime = manIntTime;
    readSpectrometer(); // read to dim0
    prevSatN = satN;
    prevIntTime = intTime;
  }
 
  Serial.print(String(panVal) + "," + String(tiltVal) + "," + String(darkLight) + "," + String(prevIntTime) + "," + String(prevSatN) );

  for (int i = 0; i < nSites; i+=boxcar){
    int tSum = 0;
    for(int j = 0; j < boxcar; j++)
      if(i+j <= nSites)
          tSum += data[i+j][dataLoc];
    Serial.print("," + String(tSum));
  }
  Serial.print("\n");
  delay(1);
 
}


void pan(long pv){
  //stepper_pan.enableOutputs();
  stepper_pan.runToNewPosition(pv);
  //stepper_pan.disableOutputs();
}

void tilt(long tv){
  //stepper_tilt.enableOutputs();
  stepper_tilt.runToNewPosition(tv);
  //stepper_tilt.disableOutputs();
}

void darkMeasure(){
      //-----------Return to zero point------------
      //panVal = 0;
      //pan(0);
      //tiltVal = 0;
      tilt(0);
      darkLight = 0; // dark measurement
      long tl = manIntTime; // save existing manual value

      // first scan at microsecond
      manIntTime = minIntTime;
      radianceMeasure();
   
      for (long i = intStep; i <= maxIntTime; i*= 2){
        manIntTime = i;
        radianceMeasure();
      }

      manIntTime = tl;// restore manIntTime
      darkLight = 1; // light measurement
}

void startMeasure(){
      //-----------Return to zero point------------
      //panVal = 0;
      pan(0);
      //tiltVal = 512;
      tilt(512);
      darkLight = 2; // dark measurement
      radianceMeasure();
      darkLight = 1; // light measurement
}


void loop() {

  
  String arg = Serial.readString();

  if (arg != NULL){

    // manually set integration time
    if(arg.startsWith("t") == true){
      arg.replace("t", "");
      manIntTime = (long) arg.toFloat();
      if(manIntTime > maxIntTime)
          manIntTime = maxIntTime;
      Serial.println("int. time: " + String(manIntTime) + "ms");

    // manual pan
    } else if(arg.startsWith("p") == true){
      arg.replace("p", "");
      stepper_pan.enableOutputs();
      pan((int) arg.toFloat());
      stepper_pan.disableOutputs();
      Serial.println("pan: " + String((int) arg.toFloat()));
      delay(5);

    // manual tilt
    } else if(arg.startsWith("l") == true){
      arg.replace("l", "");
      stepper_tilt.enableOutputs();
      tilt((int) arg.toFloat());
      stepper_tilt.disableOutputs();
      Serial.println("tilt: " + String((int) arg.toFloat()));
      delay(5);

    // Manual spec measure
    } else if(arg.startsWith("r") == true){ // radiance

      radianceMeasure();

    // -------------  hyperspec --------------
    } else if(arg.startsWith("h") == true){
      arg.replace("h", "");
      int i= 0;
      while(arg.indexOf(",") != -1){
        hyperVals[i] = long(arg.substring(0,arg.indexOf(",")).toFloat()); //toInt for int, atol for long
        i++;
        arg = arg.substring(arg.indexOf(",")+1);
      }

   

      Serial.println("h," + String(unitNumber) + "," + String(hyperVals[0]) + "," + String(hyperVals[1]) + "," + String(hyperVals[2]) + "," + String(hyperVals[3]) + "," + String(hyperVals[4]) + "," + String(hyperVals[5]) + "," + String(hyperVals[6]) + "," + String(hyperVals[7]) + "," + String(hyperVals[8]) );
      delay(5);
      //arg = Serial.readString();
      
      maxIntTime = (long) hyperVals[6];
      boxcar = (int) hyperVals[7];
      darkRepeat = (long) hyperVals[8];

      stepper_pan.enableOutputs();
      stepper_tilt.enableOutputs();
      //panVal = 0;
      pan(0);
      tilt(0);

      //-----------Dark Measure at start------------
      startMeasure();
      darkMeasure();

      unsigned long cDR = millis(); // current time dark repeat
      unsigned long eDR = cDR + darkRepeat; // end time dark repeat
      
      for(tiltVal = hyperVals[3]; tiltVal <= hyperVals[4]; tiltVal += hyperVals[5]){
        
        //----------repeat dark measurement based on timeout-----------
        cDR = millis();
        if(cDR >= eDR){
            darkMeasure();
            //tilt(tiltVal);
            cDR = millis();
            eDR = cDR + darkRepeat;
        }

        tilt(tiltVal);
        pan(hyperVals[0]-10);// overshoot - pan left a bit to use up excess in gears

        //--------reduce measurement frequency at high elevations
        int panShift = 1;
        int panStart = 0;
        for(int i=0; i<4; i++)
        if(tiltVal >= panoSteps[i]){
            panShift = panoSpaces[i];
            panStart = panoSpaces[i]/2;
        }
        panShift *= hyperVals[2];
        panStart *= hyperVals[2];
            
        /*
        if(panVal >= hyperVals[1]){
          //----------pan from right to left-----------
          for(panVal = hyperVals[1]-panStart; panVal >= hyperVals[0]; panVal -= panShift){ 
            pan(panVal);
            radianceMeasure();
          }
        } else {
        */
          //----------pan from left to right-----------
          for(panVal = hyperVals[0]+panStart; panVal <= hyperVals[1]; panVal += panShift){ 
            pan(panVal);
            radianceMeasure();
          }
        //}

      }


      //-----------Dark Measure at end------------
      pan(0);
      darkMeasure();

      stepper_pan.disableOutputs();
      stepper_tilt.disableOutputs();

      Serial.println("x"); // signal end
//      Serial.println("\n"); // signal end
    }
  }

  delay(10);  
     
  


}
```
