## Supplementary material for "Hyperspectral open source imaging system": HOSI supplementary files including code and 3D printer models: Calculate_HOSI_linearisation_curve_0.6.py

```
##
##_________________________HOSI calculate linearisation coefficients_____________________________
##
## Written by: Jolyon Troscianko
## initial release: 19/06/2024
## License: GNU General Public License v3.0
##
## To use this fucntion, point the HOSI so that it's facing a stably illuminated surface
## then just run the script. Ideally do this in a dark room. It is essential that the
## illuminat is stable for the whole minute or so that this runs.
##
## Adjust the brightness of the source so that the longest exposures are in the range
## of around 200,000 to 800,000 microseconds (to ensure a nice wide dynamic range is used
##
## Once finished, check the graphs look good and R^2 values are ideally >0.999
## then copy the two coeffients from the log to the linCoefs in calibration_data.txt


import serial, time
import serial.tools.list_ports
import numpy as np
import matplotlib.pyplot as plt
from scipy.optimize import curve_fit
from scipy.ndimage import gaussian_filter


ports = serial.tools.list_ports.comports()
com_list = []
for p in ports:
	com_list.append(p.device)

print(com_list)
#serialName = '/dev/ttyUSB0'
if(len(com_list) > 0):
	serialName = com_list[0]
	ser = serial.Serial(serialName, 115200) #115200 230400
else:
	serialName = 0


baseInt = 550 # specifies the minimum hardware integration time in microseconds.  
minInt = 1000 # ignore integration times shorter than this value
maxCount = 1000 # saturation typically occurs around 1000 for arduino reads of counts from Hamamatsi chip (ADC range up to 1023)


time.sleep(3) # the serial monitor seems to need this time to work!

#-------TILT--------------
ser.write(b'l500')
print("Move to start")
serFlag = 0
while serFlag == 0: # wait for tilt to get to where it's going
	output = ser.readline()
	#print(output)
	if(output.startswith(b't')):
		serFlag = 1

# take test auto-exposure
ser.write(b'r')
print("Radiance test")
serFlag = 0
while serFlag == 0: # wait for tilt to get to where it's going
	output = ser.readline()
	#print(output)
	if(output.startswith(b'0')):
		serFlag = 1

output = output.decode('utf_8')
output = output.split(',')
expAuto = int(output[3])
print("Auto exposure: " + str(expAuto))

output = output[5:] # remove first 5 values
#outputInts = [int.from_bytes(output, byteorder='little') for output in output] # convert to int
outputInts = [int(output) for output in output]

# peak spec value
maxVal = max(outputInts)

# Find the index of the maximum value
maxIndex = outputInts.index(maxVal)
print("Max photosite: " + str(maxIndex) + " MaxVal:" + str(maxVal))

exps = [0.005, 0.006, 0.007, 0.008, 0.009, 0.01, 0.02, 0.05, 0.1, 0.2, 0.3, 0.4, 0.5, 0.6, 0.7, 0.8, 0.9, 1.0, 1.1, 1.2, 1.4, 1.6]
lightData = np.array([])
darkData = np.array([])
intTimes = np.array([])
satData = np.array([])


for i in exps:
	if int(expAuto * i) > minInt:
		ts = "t" + str(int(expAuto * i))
		#print(ts)
		ser.write(str.encode( ts ))
		serFlag = 0
		while serFlag == 0: # wait for tilt to get to where it's going
			output = ser.readline()
			#print(output)
			if(output.startswith(b'i')): # int time
				serFlag = 1
		ser.write(b'r')
		serFlag = 0
		while serFlag == 0: # wait for tilt to get to where it's going
			output = ser.readline()
			#print(output)
			if(output.startswith(b'0')):
				serFlag = 1
		output = output.decode('utf_8')
		output = output.split(',')
		output = output[5:] # remove first 5 values
		outputInts = [float(output) for output in output]
		outputSmoothed = gaussian_filter(outputInts, 3)

##		print("Integration Time: " + str(int(expAuto * i)) + " Peak: " + str(outputInts[maxIndex]))
		satData = np.append(satData, outputInts[maxIndex])
		print("Integration Time: " + str(int(expAuto * i)) + " Peak: " + str(outputSmoothed[maxIndex]))
		lightData = np.append(lightData, outputSmoothed[maxIndex])
		intTimes = np.append(intTimes, int(expAuto * i))


#--------CLOSE---------
time.sleep(1)
ser.write(b'l0')
print("Close")
serFlag = 0
while serFlag == 0: # wait for tilt to get to where it's going
	output = ser.readline()
	#print(output)
	if(output.startswith(b't')):
		serFlag = 1


for i in exps:
	if int(expAuto * i) > minInt:
		ts = "t" + str(int(expAuto * i))
		#print(ts)
		ser.write(str.encode( ts ))
		serFlag = 0
		while serFlag == 0: # wait for tilt to get to where it's going
			output = ser.readline()
			#print(output)
			if(output.startswith(b'i')): # int time
				serFlag = 1
		ser.write(b'r')
		serFlag = 0
		while serFlag == 0: # wait for tilt to get to where it's going
			output = ser.readline()
			#print(output)
			if(output.startswith(b'0')):
				serFlag = 1
		output = output.decode('utf_8')
		output = output.split(',')
		output = output[5:] # remove first 5 values
		outputInts = [float(output) for output in output]
		outputSmoothed = gaussian_filter(outputInts, 3)
##		print("Integration Time: " + str(int(expAuto * i)) + " ave dark: " + str(np.mean(outputInts)))
##		darkData = np.append(darkData, np.mean(outputInts))
		print("Integration Time: " + str(int(expAuto * i)) + " Peak: " + str(outputSmoothed[maxIndex]))
		darkData = np.append(darkData, outputSmoothed[maxIndex])


## Remove over-exposed values
mask = satData <= maxCount

# Apply the mask to arrays
darkData = darkData[mask]
intTimes = intTimes[mask]
lightData = lightData[mask]


lightData = lightData - darkData


## Remove negative values
mask = lightData > 0

# Apply the mask to arrays
darkData = darkData[mask]
intTimes = intTimes[mask]
lightData = lightData[mask]


scaledIntTimes = ((intTimes+baseInt)/(max(intTimes)+baseInt))*max(lightData)

combined = np.column_stack((intTimes, lightData, scaledIntTimes))

# Print each row
print("intTimes, lightData, scaledIntTimes (linear)")
for row in combined:
    print(row)


def custom_equation(x, a, b):
	#return a * np.log((x+1) * b)
	#return a*x+b
	return np.exp(a*np.log(x)+b)


# Perform the curve fitting
initial_guess = [1.0, 0.0]  # Initial guess for the coefficients
params, params_covariance = curve_fit(custom_equation, lightData, scaledIntTimes, p0=initial_guess)

# Print the fitted coefficients
print("Fitted coefficients:")
print(params)


# Calculate the residuals and R^2
y_fit = custom_equation(lightData, *params)
residuals = scaledIntTimes - y_fit
ss_res = np.sum(residuals**2)
ss_tot = np.sum((scaledIntTimes - np.mean(scaledIntTimes))**2)
r_squared = 1 - (ss_res / ss_tot)

print(f"R^2: {r_squared}")

# Plot the original data and the fitted curve
plt.figure(figsize=(8, 6))
plt.xscale('log')
plt.yscale('log')
plt.scatter(lightData, scaledIntTimes, label='Data', color='red')
plt.plot(lightData, custom_equation(lightData, *params), label='Fitted curve', color='blue')
plt.xlabel('Observed Counts')
plt.ylabel('Expected Linear Counts')
plt.legend()
plt.show()
```
