## Supplementary material for "Hyperspectral open source imaging system": HOSI supplementary files including code and 3D printer models: Calculate_HOSI_spectral_sensitivity_0.2.py

```
##
##_________________________HOSI calculate spectral sensitivity_____________________________
##
## Written by: Jolyon Troscianko
## initial release: 19/06/2024
## License: GNU General Public License v3.0
##
## 
## 
## To use this script you must already have calculated the linearidsation coefficients
## (see other script), and inputed the wavelength coefficients that come with the chip.
## 
## Using the HOSI, take an image of something with a known (calibrated) radiance. e.g.
## a diffuse white standard illuminated with a broadband light source, measured
## using a calibrated spectroradiometer. Make sure you don't use spectral compression
## i.e. set the boxcar to 1 in the HOSI GUI.
## 
## You need to extract the raw light count data from the HOSI output csv file, and from this 
## subtract the dark count data that use the same integration time. These are measured both
## before and after each scan (and intermittently for long scans). intput these raw count
## data below together with known calibration spectra and run the script. if it looks good
## then copy the comma-delineated line of spectral sensitivities to the radSens line
## in calibration_data.csv
## 
## Make sure the unit number in calibration_data.csv matches the unit number you gave to
## the Arduino when writing the firmware. I recommend editing calibraiton_data.txt
## 
## 
## 


import numpy as np
import matplotlib.pyplot as plt
from scipy.interpolate import interp1d
from scipy.ndimage import gaussian_filter


#________________UNIT_SPECIFIC DATA___________________
# you must alter each of the following unit-specific data:

wavlCoefs = np.array([297.308340799761,2.72E+00,-1.15E-03,-8.03E-06,1.24E-08,-1.48E-12])
linCoefs = np.array([0.91106809,0.58315014])
# counts are average counts minus dark measurement, plus integration time in microseconds. Note that calibration should ideally be done with boxcar=1 (i.e. no downsampling of spectra)
counts = np.array([10.043,10.214,8.900,9.414,11.514,12.743,17.400,24.700,32.229,45.271,60.757,74.457,83.971,91.514,97.929,102.057,104.586,106.786,115.086,109.543,122.600,126.957,130.200,143.229,161.686,173.886,177.086,194.586,210.071,238.543,257.214,277.814,288.957,296.000,305.400,306.000,312.514,308.400,309.886,306.200,305.557,307.371,311.971,312.200,317.571,326.200,336.543,346.943,366.514,386.843,418.557,445.586,472.400,499.229,529.600,565.643,616.571,666.386,702.400,703.557,690.471,656.229,629.286,615.900,609.743,596.014,578.929,568.700,561.471,547.729,523.143,505.143,491.171,484.900,481.000,480.557,476.400,478.543,482.057,483.871,481.943,492.143,500.186,506.543,514.286,518.286,513.500,521.586,516.143,515.843,512.343,505.029,505.643,508.529,504.729,505.557,500.871,510.800,506.829,507.700,511.443,509.000,514.371,513.271,515.414,517.943,520.471,521.414,529.686,525.386,522.557,527.086,524.200,521.886,527.800,525.971,525.600,524.886,519.771,529.971,533.200,533.886,535.800,536.071,540.029,544.929,531.443,533.700,525.843,521.214,522.900,513.657,504.871,490.743,487.929,492.914,484.000,483.600,478.757,470.886,463.557,455.314,449.414,447.171,446.843,442.286,436.271,438.000,433.800,438.457,431.800,430.086,426.714,433.357,432.814,432.571,434.929,431.543,424.400,416.800,407.100,395.214,387.129,375.200,373.857,367.471,361.700,363.957,362.200,365.929,371.929,372.729,364.400,358.957,352.400,342.629,338.300,335.300,338.400,346.543,348.600,347.486,345.829,344.529,343.929,333.729,323.800,321.714,317.371,316.014,316.571,302.829,296.886,294.214,298.457,291.000,298.657,314.700,317.800,330.857,324.386,316.843,308.857,278.429,262.400,239.800,234.100,223.329,213.057,212.529,211.986,219.343,206.400,207.014,207.657,212.986,217.386,204.057,205.114,206.114,202.243,197.471,194.586,201.129,198.157,200.586,186.371,184.729,188.929,194.314,204.571,208.457,235.643,287.986,353.171,453.229,568.814,666.143,728.586,751.186,727.786,660.871,583.157,490.629,398.543,340.743,306.243,277.386,245.486,227.143,214.814,192.471,173.743,160.057,143.129,133.686,118.886,116.886,109.543,112.457,102.557,107.671,107.943,105.671,102.143,108.929,107.100,106.943,109.829,117.729,112.643,119.800,122.486,126.900,136.300,155.100,175.957,197.343,226.171,256.986,296.614,337.586,360.943,368.629,369.743,346.929,331.386,303.129])
intTime = 51200
# Spectroradiometer wavelengths and intensity data
wavelengthsCalibrated = np.array([230,231,232,233,234,235,236,237,238,239,240,241,242,243,244,245,246,247,248,249,250,251,252,253,254,255,256,257,258,259,260,261,262,263,264,265,266,267,268,269,270,271,272,273,274,275,276,277,278,279,280,281,282,283,284,285,286,287,288,289,290,291,292,293,294,295,296,297,298,299,300,301,302,303,304,305,306,307,308,309,310,311,312,313,314,315,316,317,318,319,320,321,322,323,324,325,326,327,328,329,330,331,332,333,334,335,336,337,338,339,340,341,342,343,344,345,346,347,348,349,350,351,352,353,354,355,356,357,358,359,360,361,362,363,364,365,366,367,368,369,370,371,372,373,374,375,376,377,378,379,380,381,382,383,384,385,386,387,388,389,390,391,392,393,394,395,396,397,398,399,400,401,402,403,404,405,406,407,408,409,410,411,412,413,414,415,416,417,418,419,420,421,422,423,424,425,426,427,428,429,430,431,432,433,434,435,436,437,438,439,440,441,442,443,444,445,446,447,448,449,450,451,452,453,454,455,456,457,458,459,460,461,462,463,464,465,466,467,468,469,470,471,472,473,474,475,476,477,478,479,480,481,482,483,484,485,486,487,488,489,490,491,492,493,494,495,496,497,498,499,500,501,502,503,504,505,506,507,508,509,510,511,512,513,514,515,516,517,518,519,520,521,522,523,524,525,526,527,528,529,530,531,532,533,534,535,536,537,538,539,540,541,542,543,544,545,546,547,548,549,550,551,552,553,554,555,556,557,558,559,560,561,562,563,564,565,566,567,568,569,570,571,572,573,574,575,576,577,578,579,580,581,582,583,584,585,586,587,588,589,590,591,592,593,594,595,596,597,598,599,600,601,602,603,604,605,606,607,608,609,610,611,612,613,614,615,616,617,618,619,620,621,622,623,624,625,626,627,628,629,630,631,632,633,634,635,636,637,638,639,640,641,642,643,644,645,646,647,648,649,650,651,652,653,654,655,656,657,658,659,660,661,662,663,664,665,666,667,668,669,670,671,672,673,674,675,676,677,678,679,680,681,682,683,684,685,686,687,688,689,690,691,692,693,694,695,696,697,698,699,700,701,702,703,704,705,706,707,708,709,710,711,712,713,714,715,716,717,718,719,720,721,722,723,724,725,726,727,728,729,730,731,732,733,734,735,736,737,738,739,740,741,742,743,744,745,746,747,748,749,750,751,752,753,754,755,756,757,758,759,760,761,762,763,764,765,766,767,768,769,770,771,772,773,774,775,776,777,778,779,780,781,782,783,784,785,786,787,788,789,790,791,792,793,794,795,796,797,798,799,800,801,802,803,804,805,806,807,808,809,810,811,812,813,814,815,816,817,818,819,820,821,822,823,824,825,826,827,828,829,830,831,832,833,834,835,836,837,838,839,840,841,842,843,844,845,846,847,848,849,850,851,852,853,854,855,856,857,858,859,860,861,862,863,864,865,866,867,868,869,870,871,872,873,874,875,876,877,878,879,880,881,882,883,884,885,886,887,888,889,890,891,892,893,894,895,896,897,898,899,900,901,902,903,904,905,906,907,908,909,910,911,912,913,914,915,916,917,918,919,920,921,922,923,924,925,926,927,928,929,930,931,932,933,934,935,936,937,938,939,940,941,942,943,944,945,946,947,948,949,950,951,952,953,954,955,956,957,958,959,960,961,962,963,964,965,966,967,968,969,970,971,972,973,974,975,976,977,978,979,980,981,982,983,984,985,986,987,988,989,990,991,992,993,994,995,996,997,998,999,1000])
intensitiesCalibrated = np.array([0.00000429667,0.000000964177,0.00000291383,0.000000463614,0.00000455703,0.00000513648,0.00000114807,0,0.00000131206,0.000000221669,0.00000102539,0.000000381009,0.00000114129,0.00000191642,0.00000201349,0,0.00000150739,0.0000020896,0.00000156959,0.0000022669,0.00000197287,0.00000255764,0.00000285747,0.00000566012,0.00000274448,0.00000408334,0.0000025924,0.00000218251,0.00000406137,0.00000347706,0.00000469114,0.00000373877,0.00000381733,0.00000567131,0.00000702561,0.00000460726,0.00000534604,0.00000451433,0.00000691736,0.00000614872,0.00000553216,0.00000592173,0.00000461776,0.00000767887,0.00000555379,0.00000676157,0.00000628539,0.00000624743,0.00000726657,0.00000757364,0.0000070372,0.00000708044,0.00000732096,0.00000760869,0.00000784476,0.00000790313,0.00000799872,0.00000864528,0.00000768502,0.00000949971,0.00000864557,0.00000940678,0.00000901407,0.00000911325,0.0000103186,0.000010131,0.0000101773,0.0000105037,0.0000109814,0.0000106177,0.0000116039,0.000011772,0.0000116829,0.0000117869,0.0000124714,0.000012366,0.0000134926,0.0000134867,0.0000140611,0.0000143649,0.0000142929,0.0000146825,0.0000151644,0.0000154548,0.0000152396,0.0000160032,0.0000154868,0.0000163941,0.0000167673,0.0000173757,0.0000178843,0.0000178216,0.0000197728,0.000019967,0.0000203955,0.0000208761,0.000021671,0.0000224299,0.000022631,0.000024116,0.0000263218,0.0000286939,0.0000288638,0.0000306669,0.0000330754,0.0000347488,0.00003667,0.0000372572,0.0000400516,0.0000444734,0.0000485397,0.000052276,0.0000553425,0.0000594058,0.0000616994,0.0000636487,0.0000666212,0.0000697502,0.000073319,0.0000751295,0.0000777159,0.000079481,0.0000810932,0.0000824987,0.0000835089,0.0000850489,0.0000869847,0.0000889783,0.0000906129,0.0000938453,0.0000973474,0.000100655,0.000104858,0.000108539,0.000112916,0.000116152,0.000120464,0.000124966,0.000127102,0.000129209,0.000130784,0.000131758,0.000134063,0.000134316,0.000134381,0.000137311,0.000141207,0.00014261,0.000145368,0.000147806,0.000149466,0.000150408,0.000152547,0.000153339,0.000154776,0.000159084,0.000161999,0.000164144,0.000165456,0.00016822,0.000171361,0.000174175,0.000177292,0.000178828,0.000181436,0.000183515,0.000187395,0.000192127,0.00019734,0.000201003,0.000202417,0.000203911,0.000206262,0.000207536,0.000207967,0.000210222,0.000211591,0.000212463,0.000215947,0.000219118,0.00022074,0.00022272,0.00022512,0.000226767,0.000230292,0.000230572,0.00023087,0.000233185,0.000236379,0.000238589,0.00024236,0.000243358,0.000244693,0.000244435,0.000243515,0.000244067,0.000244401,0.000246969,0.000244871,0.000246013,0.000246115,0.00024864,0.00025012,0.00025347,0.000256062,0.000258033,0.000260839,0.000263829,0.000265629,0.000269641,0.000271354,0.000271322,0.000272701,0.000274784,0.000275666,0.000276449,0.000277138,0.000281819,0.000288656,0.000298153,0.000308688,0.000315544,0.000316406,0.00031294,0.000309487,0.000307146,0.000308647,0.00031492,0.000319673,0.000323347,0.000328623,0.000334956,0.000344353,0.000353168,0.000360384,0.000367894,0.000376794,0.000391604,0.000405506,0.000413408,0.000411456,0.0004045,0.000397144,0.000392625,0.000385009,0.000373414,0.000361008,0.000351451,0.000345207,0.000346391,0.000351476,0.000361054,0.000369039,0.000371628,0.000369406,0.000363363,0.000355527,0.000350116,0.000346984,0.000347482,0.000351175,0.000359165,0.000365896,0.000369305,0.000369306,0.000366878,0.00035935,0.000353547,0.000349794,0.000348125,0.000346477,0.00034575,0.000344759,0.000346057,0.000347598,0.000348796,0.000348349,0.00034838,0.000348541,0.000349737,0.000351836,0.00035367,0.00035364,0.000354869,0.000356017,0.000355829,0.000357277,0.000357571,0.000358216,0.000359728,0.000361725,0.000363651,0.000363773,0.000365271,0.00036683,0.000367902,0.000368009,0.000367242,0.000369122,0.000370842,0.00037118,0.000371853,0.000373173,0.00037405,0.000374196,0.000376194,0.00037589,0.000377187,0.000377754,0.000378398,0.000379536,0.000380155,0.000381591,0.000384153,0.000384458,0.000386138,0.000385735,0.000387103,0.000386923,0.000387496,0.00038801,0.000388738,0.00038996,0.000391393,0.000392141,0.0003924,0.000393191,0.000393661,0.000394237,0.000394936,0.000396831,0.000396036,0.000397637,0.000397619,0.000398178,0.000398652,0.000400384,0.000400494,0.000402173,0.000403633,0.000405376,0.000406597,0.000406766,0.000408621,0.000409465,0.000410269,0.000411272,0.000412827,0.000411819,0.00041382,0.000413961,0.000416271,0.000416342,0.000419082,0.000420239,0.000422509,0.000421312,0.000422672,0.000423641,0.000424466,0.000425394,0.000425624,0.000426667,0.000426769,0.000427289,0.000426273,0.000425308,0.000425717,0.000426243,0.000425749,0.000425399,0.00042661,0.000424914,0.000426859,0.000427108,0.000427029,0.000428179,0.000427962,0.000430268,0.000431524,0.000431953,0.000433306,0.000434013,0.000435234,0.0004384,0.000441124,0.000443257,0.000445202,0.000447379,0.000450337,0.000452232,0.000455133,0.000456188,0.000455948,0.000456649,0.000454795,0.000454579,0.000454234,0.000452306,0.000453128,0.000453339,0.000454134,0.000453769,0.000454432,0.000455882,0.000455483,0.0004552,0.000454136,0.000452264,0.000450234,0.00045081,0.000449656,0.000449545,0.000449906,0.000448655,0.000451796,0.000453482,0.000456193,0.000459615,0.000462805,0.00046542,0.000465691,0.000462982,0.000461783,0.000457083,0.000454446,0.000453278,0.000452939,0.000450775,0.000452744,0.00045383,0.000455584,0.000458633,0.000458627,0.000460171,0.000463179,0.000465005,0.000469122,0.000470952,0.000473899,0.000476096,0.000476892,0.000477983,0.000479923,0.000480627,0.000483118,0.000484232,0.000483055,0.000485488,0.000485842,0.000488333,0.000491651,0.000498284,0.000508653,0.000516409,0.000521284,0.000523401,0.000524071,0.000524869,0.000529845,0.000532715,0.000534988,0.000532772,0.000526539,0.000518292,0.000509996,0.000504508,0.000498673,0.000495466,0.000492985,0.000490667,0.000489792,0.000487941,0.00048541,0.000484255,0.000484386,0.000483045,0.000485781,0.000489437,0.000495743,0.0005033,0.000515311,0.000528289,0.000539576,0.000548016,0.000553323,0.000550658,0.000544092,0.00053607,0.000527335,0.000518327,0.000509301,0.000504869,0.000502933,0.00050041,0.00050151,0.000504805,0.000511566,0.000523116,0.00052912,0.000543835,0.000557654,0.000572667,0.000584961,0.000592169,0.000589229,0.000577982,0.000563412,0.000554679,0.000550798,0.000555329,0.000560782,0.00056611,0.000569647,0.000569549,0.000564828,0.000558714,0.00055477,0.000550333,0.000554053,0.0005553,0.000557529,0.000558977,0.000556826,0.000553304,0.000550795,0.000552103,0.000549721,0.000551322,0.000553223,0.000558771,0.000576757,0.000615988,0.000674404,0.000751224,0.000810062,0.000815641,0.000768785,0.000692284,0.00062574,0.000583525,0.000558452,0.000543446,0.000534796,0.000528056,0.000518831,0.000512073,0.00050538,0.0005003,0.000497126,0.000495241,0.0004961,0.000496908,0.000497837,0.00049941,0.000499298,0.000502064,0.000513368,0.000528957,0.000546053,0.000561531,0.000565855,0.000556098,0.000545923,0.000535712,0.000533096,0.000536842,0.000547568,0.000557272,0.000563606,0.000562989,0.000563674,0.00055584,0.000546712,0.000539911,0.000535969,0.000537415,0.000540951,0.000543815,0.000550839,0.000556489,0.000561067,0.000566407,0.000574994,0.000585839,0.000600804,0.000623104,0.000658942,0.000722318,0.000860901,0.00115207,0.00167261,0.00243668,0.00319343,0.00366291,0.00367405,0.00331808,0.00287328,0.00250164,0.00218171,0.00187172,0.00157498,0.00138667,0.00132505,0.00136297,0.00139074,0.00134782,0.00123016,0.00108368,0.000979519,0.000926593,0.000905972,0.000876804,0.000815395,0.000737436,0.000659755,0.000596631,0.000553582,0.000527594,0.00051373,0.000500786,0.000495442,0.000492573,0.000490707,0.000489138,0.000490313,0.000491329,0.00049426,0.000496752,0.000503208,0.000506758,0.000514659,0.00052197,0.000531875,0.000542382,0.000556604,0.000574957,0.00059185,0.000613613,0.000635892,0.000658862,0.000682111,0.000711188,0.000746936,0.000784669,0.000836538,0.000918642,0.00107546,0.00136863,0.00187751,0.00255415,0.00323491,0.00380547,0.00406419,0.00396712,0.00358202,0.00300571,0.00241554,0.00194,0.00162543,0.00145229,0.00141443,0.00147337,0.00159031,0.00173361,0.00184726,0.00190802,0.00188519,0.00177255,0.00163797,0.00153524,0.00152427,0.00161179,0.00176385,0.00191853,0.00202653,0.00204678,0.00194196,0.00175859,0.00152764,0.00135461,0.00129397,0.00137392,0.00157187,0.00183062,0.0020945,0.00233061,0.00248111,0.00250487,0.00239441,0.00217873,0.00190368,0.0016379,0.0014195,0.00124794,0.00112419,0.00103421,0.000965456,0.000914942,0.000871975,0.00085001,0.000836238,0.000841266,0.00085107,0.000875611,0.00090807,0.000949289,0.00100523,0.00105662,0.00110745,0.00116047,0.00119924,0.00122512,0.00124245,0.00123895,0.00124103,0.0012399,0.00124076,0.00124657,0.00125225,0.00125561,0.00126095,0.00125848,0.00123817,0.0012089,0.00116725,0.0011112,0.00105247,0.000994873,0.000933276,0.000885899,0.000847439,0.00081881,0.000803889,0.00079797,0.000802937,0.000802742,0.000820113,0.000843369,0.000874474,0.000910067,0.000951711,0.000993064,0.00104289,0.00112365,0.00123035,0.00136436,0.00154223,0.00172144,0.00189286,0.00204361,0.00214303,0.0022119,0.00222936,0.00215948,0.00201585,0.00183915,0.00165476,0.00156249,0.00154497,0.00161411,0.00175164,0.00191261,0.00206323,0.0021845,0.00226727,0.00226704,0.00220872,0.00207717,0.00187755,0.00166951])


#_______________Calculations_____________
##countsLin =  counts / ( linCoefs[0]*np.log((counts+1)*linCoefs[1]))
countsLin = np.exp(np.log(counts)*linCoefs[0]+linCoefs[1])
countsLinSmoothed = gaussian_filter(countsLin, 3)
photosites = np.arange(288)
wavelengths = wavlCoefs[0]+wavlCoefs[1]*photosites+wavlCoefs[2]*photosites**2+wavlCoefs[3]*photosites**3+wavlCoefs[4]*photosites**4+wavlCoefs[5]*photosites**5

# Interpolate calibration spectrum to the wavelengths of the microspec
interp_func = interp1d(wavelengthsCalibrated, intensitiesCalibrated, kind='linear', bounds_error=False, fill_value='extrapolate')
intensitiesCalibratedAligned = interp_func(wavelengths)
intensitiesCalibratedAlignedSmoothed = gaussian_filter(intensitiesCalibratedAligned, 6)

sensitivities = countsLinSmoothed/(intTime*intensitiesCalibratedAlignedSmoothed)


# Plot the original and aligned spectra
plt.figure(figsize=(10, 6))
plt.plot(wavelengths, sensitivities, label='Sensitivities', color='blue')
plt.xlabel('Wavelength (nm)')
plt.ylabel('Intensity')
plt.title('Spectrum Alignment')
plt.legend()
plt.show()

#print(sensitivities)
#sensitivitiesOutput = ','.join(map(str, sensitivities))
sensitivitiesOutput = [f"{x:.5f}" for x in sensitivities]
sensitivitiesOutput = ",".join(sensitivitiesOutput)

print(sensitivitiesOutput)
```
