## Supplementary material for "Hyperspectral open source imaging system": HOSI supplementary files including code and 3D printer models: HOSI_GUI_0.1.45.py

```
##
##_________________________HOSI GUI_____________________________
##
## Written by: Jolyon Troscianko
## initial release: 19/06/2024
## License: GNU General Public License v3.0
##
## This script creates a GUI for running a hyperspectral open source imager (HOSI) device
## designed to work on linux or Android operating systems. To use on Android, install
## the free Pydroid 3 app, and run the code from there. You'll need to install the dependencies
## using PIP in Pydroid, which are:
##
##	matplotlib
##	usb4a
##	usbserial4a
##
## Units require unit-specific calibraiton and testing. Calibration coefficients must be present
## in the calibration_data.txt file in the same directory as this script. You also need to have
## present the grid.png file, and sensitivity_data.csv that contains human spectral sensitivities
##
##
##
## When the lens is as far away from the sensor as possible, the focus distance is about 30mm
##
##


from tkinter import *
from tkinter import filedialog as fd
import numpy as np
from PIL import Image, ImageTk, ImageOps
import string, time, os, sys, math

import matplotlib.pyplot as plt
from matplotlib.backends.backend_tkagg import FigureCanvasTkAgg


platform = "lin"
if any(key for key in os.environ if key.startswith('ANDROID_')) == True:
	platform = "and"
	from usb4a import usb
	from usbserial4a import serial4a

##else if( platform.system.startswith("WINDOWS")):
##        platform = "win"

else:
	import serial
	import serial.tools.list_ports

ser = None


root = Tk()
root.geometry('600x800')
root.title('HOSI')


np.set_printoptions(suppress=False, precision=3, threshold=sys.maxsize, linewidth=sys.maxsize)

cieWav = []
chlA = []
chlB = []
nIR = []
cieX = []
cieY = []
cieZ = []
nUV = []


pixels = int(288)
baseInt = int(550) # specifies the minimum hardware integration time in microseconds. 
wavCoef = []
radSens = []
linCoefs = []
wavelength = []
wavelengthBins = []
wavelengthBoxcar = []
unitNumber = int(0)
cieXt = [0.0] * pixels
cieYt = [0.0] * pixels
cieZt = [0.0] * pixels
chlAt = [0.0] * pixels
chlBt = [0.0] * pixels
nIRt = [0.0] * pixels
nUVt = [0.0] * pixels
saveLabel = StringVar()
dataString = ""
output = ""
serialName = ""
visSystems = []

def connect():
	global ser, serialName
	if platform == 'and':
		usb_device_list = usb.get_usb_device_list()
		#print(usb_device_list)
		device_name_list = [
				device.getDeviceName() for device in usb_device_list
			    ]
		#serialName ='/dev/bus/usb/001/018'
		if(len(device_name_list) > 0):
			serialName = device_name_list[0]
			deviceName = usb.get_usb_device(serialName)

			while not usb.has_usb_permission(deviceName):
				usb.request_usb_permission(deviceName)
				time.sleep(1)

			ser =  serial4a.get_serial_port(serialName, 115200,8,'N',1,timeout=1) #115200 230400
		else:
			serialName = 0
	else:
		ports = serial.tools.list_ports.comports()
		com_list = []
		for p in ports:
			com_list.append(p.device)

		print(com_list)
		#serialName = '/dev/ttyUSB0'
		if(len(com_list) > 0):
			serialName = com_list[0]
			ser = serial.Serial(serialName, 115200) #115200 230400
		else:
			serialName = 0

connect()
print(ser)

def readLine(ta):
	ta.pop(0)
	ta.pop(0)
	ta = list(filter(None, ta))
	ta = [float(i) for i in ta]
	#print(ta)
	return ta

def unitSetup():
	global unitNumber, wavCoef, radSens, linCoefs, wavelength, wavelengthBins, wavelengthBoxcar, cieXt, cieYt, cieZt, chlAt, chlBt, nIRt, nUVt, cieWav, chlA, chlB, nIR, cieX, cieY, cieZ, nUV


	if(len(wavCoef) != pixels or len(radSens) != pixels): # only load values from file the first time the code runs (to work out unit number)
		for line in open("./calibration_data.txt"):
			row = line.split(',')
			#print(row)
			try:
				int(row[0])
			except:
				print("except (not a unit number)")
			else:
				if(int(row[0]) == unitNumber):
##					print(row)
					if(row[1] == "wavCoef"):
						wavCoef = row
						wavCoef.pop(0)
						wavCoef.pop(0)
						wavCoef = list(filter(None, wavCoef))
						wavCoef.pop(len(wavCoef)-1)
##						print("waveCoef: " + str(len(wavCoef)))
						wavCoef = [float(i) for i in wavCoef]
					if(row[1] == "radSens"):
						radSens = row
						radSens.pop(0)
						radSens.pop(0)
						radSens = list(filter(None, radSens))
						radSens.pop(len(radSens)-1)
##						print("radSens: " + str(len(radSens)))
						radSens = [float(i) for i in radSens]
##						print(radSens)
					if(row[1] == "linCoefs"):
						linCoefs = row
						linCoefs.pop(0)
						linCoefs.pop(0)
						linCoefs = list(filter(None, linCoefs))
						linCoefs.pop(len(linCoefs)-1)
##						print("linCoefs: " + str(len(linCoefs)))
						linCoefs = [float(i) for i in linCoefs]
						
		for line2 in open("./sensitivity_data.csv"):
			row = line2.split(',')
##			print(row)
			if(row[0] == "base"):
				if(row[1] == "cieWav"):
					cieWav = row
					cieWav.pop(0)
					cieWav.pop(0)
					cieWav = list(filter(None, cieWav))
					cieWav = [int(i) for i in cieWav]
	##				print(cieWav)
				elif(row[1] == "chlA"):
					chlA = readLine(row)
				elif(row[1] == "chlB"):
					chlB = readLine(row)
				elif(row[1] == "nIR"):
					nIR = readLine(row)
				elif(row[1] == "cieX"):
					cieX = readLine(row)
				elif(row[1] == "cieY"):
					cieY = readLine(row)
				elif(row[1] == "cieZ"):
					cieZ = readLine(row)
				elif(row[1] == "nUV"):
					nUV = readLine(row)
##			else:
##                                for i in range(0, len(visSystems)):
##                                        TODO - add ability to export cone-catch from other spectral sensitivities
				
				
	if(len(wavCoef) != 6 or len(radSens) != pixels or len(linCoefs) != 2):
		#settingsLabel.config(text="error - calibration data not found\nEnsure calibration_data.txtis present\nand has data for unit #" + str(unitNumber))
		#return
		updateStatus("Missing calibration data")
		print("error - calibration data not found\nEnsure calibration_data.txt is present\nand has data for unit #" + str(unitNumber))
		print("waveCoef: " + str(len(wavCoef)))
		print("radSens: " + str(len(radSens)))
		print("linCoefs: " + str(len(linCoefs)))


	wavelength = []
	for i in range(0,pixels):
		#wavelength.append(float(wavCoef[0])+float(wavCoef[1])*i+float(wavCoef[2])*i**2+float(wavCoef[3])*i**3+float(wavCoef[4])*i**4+float(wavCoef[5])*i**5)
		wavelength.append(wavCoef[0]+wavCoef[1]*i+wavCoef[2]*i**2+wavCoef[3]*i**3+wavCoef[4]*i**4+wavCoef[5]*i**5)

	#---set up wavelength bin widths array-------
	wavelengthBins = []
	for i in range(0,pixels-1):
		wavelengthBins.append(wavelength[i+1]-wavelength[i])
	wavelengthBins.append(wavelength[pixels-1]-wavelength[pixels-2])

	# creare array of wavelengths matching boxcar scale for plotting
	wavelengthBoxcar = []
	for i in range(0, pixels, boxcarN): #increment in boxcar size
		wavelengthBoxcar.append(wavCoef[0]+wavCoef[1]*i+wavCoef[2]*i**2+wavCoef[3]*i**3+wavCoef[4]*i**4+wavCoef[5]*i**5)


	#------resample spectral sensitivities at spectrometer wavelengths---------
	for i in range(0,pixels):
		for j in range(0, len(cieWav)):
			if(round(wavelength[i]) == cieWav[j]):
				cieXt[i] = cieX[j]
				cieYt[i] = cieY[j]            
				cieZt[i] = cieZ[j]
				chlAt[i] = chlA[j]
				chlBt[i] = chlB[j]            
				nIRt[i] = nIR[j]
				nUVt[i] = nUV[j]


panFrom = StringVar()
panTo = StringVar()
panResolution = StringVar()
tiltFrom = StringVar()
tiltTo = StringVar()
tiltResolution = StringVar()
maxIntTime = StringVar()
boxcarVal = StringVar()
darkRepVal = StringVar()
reflVal = StringVar()

panFrom.set("-20")
panTo.set("20")
panResolution.set("4")
tiltFrom.set("480")
tiltTo.set("520")
tiltResolution.set("4")
maxIntTime.set("2000") # max int time microseconds
boxcarVal.set("2")
darkRepVal.set("120")
tt = time.time()
reflVal.set("99")

reflFlag = 0

darkTimes = []
darkVals = []

## RGB image
imLum = []
imR = []
imG = []
imB = []
imCol = []
maxRGB = 1E-10
imSatR = []
imSatB = []

## extreme spectral range image
imI = []
imGG = []
imU = []
maxIGU = 1E-10

## NDVI image
imChlA = []
imChlB = []
imNDVI = []


## hspec image - le values in array
hspec = []
hspecPan = []
hspecTilt = []


panStart = int(0)
panStop = int(0)
pan_Res = int(0)
panDim = int(11) ## initially 11 based on other start settings
tiltStart = int(0)
tiltStop = int(0)
tilt_Res = int(0)
tiltDim = int(11)

scanningFlag = int(0)
stopFlag = int(0)
fileImportFlag = int(0);
loadPath = ""
loadLine = 0
lines = ("")


boxcarN = int(1)
focusPos = 0
preview = 1
plotImX = 100
plotImY = 100
selX = -1
selY = -1
refs = [] #100% reflectance vals
wbR = 1.0
wbG = 1.0
wbB = 1.0
wbI = 1.0
wbGG = 1.0
wbU = 1.0

def updateStatus(ts):
    statusLabel.config(text=ts)

def plotGraph(status):
	global plotImX, plotImY, wbR, wbG, wbB, wbI, wbGG, wbU
##        print("plotting")
	# normalise to max=1 and non-linearise with power function
	if len(imR) > 0:
		
		br = 100/brightnessScale.get()
		if preview <= 2:
                        ## avoid raising negative numbers to power (numpy doesn't like it)
			nImR = np.sign(imR) * ((np.abs(imR * wbR)/maxRGB)**0.42) * 255 * br
			nImG = np.sign(imG) * ((np.abs(imG * wbG)/maxRGB)**0.42) * 255 * br
			nImB = np.sign(imB) * ((np.abs(imB * wbB)/maxRGB)**0.42) * 255 * br
			
##			nImR = ((imR/maxRGB)**0.42) * 255 * br
##			nImG = ((imG/maxRGB)**0.42) * 255 * br
##			nImB = ((imB/maxRGB)**0.42) * 255 * br

			nImR = np.clip(nImR, 0, 255)
			nImG = np.clip(nImG, 0, 255)
			nImB = np.clip(nImB, 0, 255)

			if preview <= 1: ## RGB images
				imCol = np.dstack((nImR, nImG, nImB))
			else: ## saturation image

				nImR = nImG ## where saturated turn red, otherwise grey (match green)
				nImB = nImG - imSatR ## where saturated turn red
				nImG = nImG - imSatR ## where saturated turn red
				nImB = np.clip(nImB, 0, 255)
				nImB = nImB + (imSatB * 5) ## add blue to show degere of saturation across wavelengths, so magenta will be fully saturated
				nImR = np.clip(nImR, 0, 255)
				nImG = np.clip(nImG, 0, 255)
				nImB = np.clip(nImB, 0, 255)
				imCol = np.dstack((nImR, nImG, nImB))

		if preview == 3:  ## IGU (extreme spectral range image)
			nImI = np.sign(imI) * ((np.abs(imI * wbI)/maxIGU)**0.42) * 255 * br
			nImGG = np.sign(imGG) * ((np.abs(imGG * wbGG)/maxIGU)**0.42) * 255 * br
			nImU = np.sign(imU) * ((np.abs(imU * wbU)/maxIGU)**0.42) * 255 * br
			
##			nImI = ((imI/maxIGU)**0.42) * 255 * br
##			nImGG = ((imGG/maxIGU)**0.42) * 255 * br
##			nImU = ((imU/maxIGU)**0.42) * 255 * br
						
			nImI = np.clip(nImI, 0, 255)
			nImGG = np.clip(nImGG, 0, 255)
			nImU = np.clip(nImU, 0, 255)

			imCol = np.dstack((nImI, nImGG, nImU))

		if preview == 4:  ## NDVI
			nImB = imChlB * 255
			nImR = ((255-nImB)*2)* imChlA
			nImG = ((255-nImB)*2)* (1-imChlA)

			imCol = np.dstack((nImR, nImG, nImB))

		# width: plot_frame.bbox(plot)[2] height: plot_frame.bbox(plot)[3]
		
##		widgetDims = root.bbox(0, 3)
		if(plot_frame.bbox(plot)[2] < plot_frame.bbox(plot)[3]):
			plotSize = plot_frame.bbox(plot)[2]
		else: plotSize = plot_frame.bbox(plot)[3]
##		plotSize = widgetDims[2] # width
##		if(widgetDims[3] < plotSize):
##			plotSize = widgetDims[3] # use height if smaller
		plotSize -= 2 ## padding

		if(plotSize < 50):
			plotSize = 50 ## set min plot size to avoid drawing errors
			
		tImCol = imCol.astype(np.uint8)
		plotIm = Image.fromarray(tImCol, "RGB")
##		plotImt = ImageOps.contain(plotIm, (plotSize,plotSize), method=0)
		plotImt = ImageOps.contain(plotIm, (plotImX,plotImY), method=0)
		plotImResized = ImageTk.PhotoImage(plotImt)
		plot.config(image=plotImResized)
		plot.image = plotImResized

		#statusLabel.config(text=status)
	   # print(root.bbox(0, 1))

##        ax.clear()
##        ax.imshow(imCol)
##        canvas.draw()
	#global panDim, pan, tilt, tiltDim
	#ts = "Scanning " +  str(float(pan + (tilt * panDim)) / float(tiltDim * panDim) + "% done")
	else:
		
		if(serialName == 0):
			statusLabel.config(text="Disconnected")
			btStart["state"] = "disabled"
##		else:
##			statusLabel.config(text="Ready")	
		
	
def updatePlotRes(event):
        global plotImX, plotImY
        plotImX = plot_frame.bbox(plot)[2]
        plotImY = plot_frame.bbox(plot)[3]
        plotGraph("Ready")


def getSpec():
	global tt, unitNumber, imLum, imR, imG, imB, imCol, imSatR, imSatB, panStart, panStop, pan_Res, panDim, tiltDim, tiltStart, tiltStop, tilt_Res, tiltRes, scanningFlag, dataString, boxcarN, maxRGB, output, focusPos
	global imI, imU, imGG, imChlA, imChlB, imNDVI, maxIGU, hspec, hspecPan, hspecTilt, fileImportFlag, loadPath, loadLine, lines, selX, selY, wavelengthBoxcar, stopFlag
	if(scanningFlag == 0 and fileImportFlag == 0): # start scanning
		# note addition of 000 to convert max int to microseconds
		ts = "h" + str(panLeft.get()) + "," + str(panRight.get()) + "," + str(panRes.get()) + "," + str(tiltBot.get()) + "," + str(tiltTop.get()) + "," + str(tiltRes.get()) + "," + str(maxInt.get()) + "000," + str(boxcar.get()) + "," + str(darkRep.get()) + "000," # note addition of three zeros for darkRep as it's expecing milliseconds
		#print(ts)
		if(int(panRight.get()) > int(panLeft.get()) and int(tiltTop.get()) > int(tiltBot.get())): # check pan & tilt coords make sense
			boxcarN = int(boxcar.get())
			#updateStatus(ts)
			statusLabel.config(text="Starting")
			ser.write(str.encode( ts ))
			#btStart["state"] = "disabled"
			btLoad["state"] = "disabled"
			btStart["text"] = "Stop"
			print("starting")
			scanningFlag = 1
			tt = time.time()
			maxRGB = 1E-10
			maxIGU = 1E-10
		else:
			statusLabel.config(text="Invalid pan/tilt")
			return

##	specLength = math.ceil(pixels/int(boxcar.get()))
		
	serFlag = 0

	if(fileImportFlag == 1 and loadLine == 0):
##		print("loading spec")
		f=open(loadPath)
		lines=f.readlines()

	while (serFlag == 0 or fileImportFlag == 1):
		if(stopFlag == 1):
			output = "x"
			dataString += output
		else:
			if(fileImportFlag == 0):
				output = ser.readline()
				try:
					output = output.decode('utf-8', 'replace')
					dataString += output
				except:
					output = "0"
					print("Error reading line")
			else:
				output = lines[loadLine]
	##			print(output)
	##			print(loadLine)
				loadLine +=1
			
			### load file & read first line

		if(output.startswith('x')):
##			print("a")
			statusLabel.config(text="Done")
			## loop to add hspec le values
			hspec = np.nan_to_num(hspec)# convert NaNs to zeros
			if(fileImportFlag == 0):
##				print("b")
				dataString2 = "le values\npan,tilt,wavelength\n,"
				for i in range(0, len(wavelengthBoxcar)):
					dataString2 += "," + str(int(wavelengthBoxcar[i]))
##				print("c") ## FAILS after this point... WHY!?
##				dataString2 += str(hspec)
				for i in range(0, len(hspecTilt)):
					for j in range(0, len(hspecPan)):
						dataString2 += '\n' + str(hspecPan[j]) + ',' + str(hspecTilt[i]) + ','
						dataString3 = str(hspec[i, j])
##						dataString3 = dataString3.replace('\n', ',')
						dataString3 = dataString3.replace(' ', ',')
						dataString3 = dataString3.replace('[', '')
						dataString3 = dataString3.replace(']', '')
						dataString3 = dataString3.replace('0.000e+00', '0')

						dataString2 += dataString3
##						for k in range(0, hspec.shape[2]):
##							try:
##								if(hspec[i, j, k] >= 0):
##									ts = f'{hspec[i, j, k]:.3e}'							
##							except:
##								print("Failed to parse output line; pan: " + str(hspecPan[j]) + " tilt: " + str(hspecTilt[i]))
##							dataString2 += ',' + ts
			
##			print("d")
			#-------save output file--------
			if(fileImportFlag == 0):
				ts = saveLabel.get()
				t = time.localtime()
				ct = "./scans/" + str(t.tm_year) + "-" + str(t.tm_mon) + "-" + str(t.tm_mday) + "_" + time.strftime("%H-%M-%S", t) + "_" + ts
				ctt = ct + ".csv"
##				print("e")
				file_object = open(ctt, 'a')
				file_object.write(dataString)
				file_object.write(dataString2)
				file_object.close()
##				print("f")
				statusLabel.config(text="Ready")
				plotGraph("")
				ctf = ct + "_sRGB.png"
	##            plt.imsave(ctf, imCol)

				#save image
				nImR = ((imR/maxRGB)**0.42) * 255
				nImG = ((imG/maxRGB)**0.42) * 255
				nImB = ((imB/maxRGB)**0.42) * 255
				imCol = np.dstack((nImR, nImG, nImB))
				
				tImCol = imCol.astype(np.uint8)
				img = Image.fromarray(tImCol, "RGB")
				#img.show()
				img.save(ctf)
##				print("g")
			
			dataString = ""
			scanningFlag = 0
			btStart["text"] = "Start"
			btStart["state"] = "active"
			btLoad["state"] = "active"
			focusPos = 0 # reset focus position in case it was previously up
			#statusLabel.config(text="Ready")
			serFlag = 1
			fileImportFlag = 0
			loadLine = 0
			selX = -1
			selY = -1 # reset these values to clear reflectance too
			stopFlag = 0
			print("h - done")
			return
		#output = output.decode('utf-8')
		output = output.split(',')
##		specLength = math.ceil(pixels/boxcarN)
		if(output[0] == 'h'):
			unitNumber = int(output[1])
			#print("unit: " + str(unitNumber))
			
			panStart = int(output[2])
			panFrom.set(str(panStart))
			panStop = int(output[3])
			panTo.set(str(panStop))
			pan_Res = int(output[4])
			panResolution.set(str(pan_Res))
			panDim = int(0)
			
			boxcarN = int(output[9])
			boxcarVal.set(str(boxcarN))
			specLength = math.ceil(pixels/boxcarN)
##			specLength = math.ceil(pixels/boxcarN)
			if(reflFlag == 1):
				clearRefl()
			unitSetup()

			tiltStart = int(output[5])
			tiltFrom.set(str(tiltStart))
			tiltStop = int(output[6])
			tiltTo.set(str(tiltStop))			
			tilt_Res = int(output[7])
			
			tiltResolution.set(str(tilt_Res))
			panDim = int(1+(panStop-panStart)/pan_Res)
			tiltDim = int(1+(tiltStop-tiltStart)/tilt_Res)
			print("Hyperspec " + str(panDim) + " by " + str(tiltDim))
			imLum = np.zeros([tiltDim, panDim])
			imR = np.zeros([tiltDim, panDim])
			imG = np.zeros([tiltDim, panDim])            
			imB = np.zeros([tiltDim, panDim])
			imCol = np.zeros([tiltDim, panDim, 3])
			imSatR = np.zeros([tiltDim, panDim])
			imSatB = np.zeros([tiltDim, panDim])

			imI = np.zeros([tiltDim, panDim])
			imGG = np.zeros([tiltDim, panDim]) 
			imU = np.zeros([tiltDim, panDim])
			
			imChlA = np.zeros([tiltDim, panDim])            
			imChlB = np.zeros([tiltDim, panDim])
			imNDVI = np.zeros([tiltDim, panDim, 3])
			
			hspec = np.zeros([tiltDim, panDim, specLength])
			hspecPan = np.zeros([panDim])
			hspecTilt = np.zeros([tiltDim])


		if(len(output) == hspec.shape[2]+5):
			if(int(output[2]) == 0 or int(output[2]) == 1 or int(output[2]) == 2 ):
				#time.sleep(0.01)
				#processSpec()
				if(preview > 0): ## live update
						root.after(1, processSpec) # 10 seems to work
				else:
						processSpec()
				#statusLabel.config(text="Running")
				return
		

def processSpec():
	global tt, darkTimes, darkVals, panStart, panStop, pan_Res, panDim, tiltDim, tiltStart, tiltStop, tilt_Res, tiltRes, linCoefs,  wavelength, wavelengthBins, maxRGB, boxcarN, output, maxIGU, hspec
	boxedLength = len(output) - 5
	updateDuringDark = 0
	if(int(output[2]) == 0): # start or restart dark measurement
		#if(int(output[3]) <= minIntTime): # start of new dark measurement when int time = 1 (might be better to comapre to previous value)
		if(len(darkTimes) == 0 or int(output[3]) < darkTimes[len(darkTimes)-1]):
			darkTimes = []
			darkVals = []
			#print("new dark measurements")

		if int(output[3]) > 100000:
				updateDuringDark = 1
		darkTimes.append(int(output[3]))
		#for i in range(0,pixels):
		for i in range(5,boxedLength+5):
			darkVals.append(output[i])

		  

	if(int(output[2]) == 1): # light measurement
		for j in range(0, len(darkTimes)): # search through dark integration times
			tempTime = int(output[3])
##            if(tempTime < 0):
##                    tempTime = 1
			if(tempTime == darkTimes[j]): # found corresponding integration time, microsecond values assumed to be equal to 1ms
				#print("int " + row[3])

			   #-----------calculate radiance-----------------
				lum = 0.0
				intTime = float(output[3])
##                if(intTime < 0):
##                        intTime = float(intTime/-1000.0)
				#if(intTime < 0):
				#        intTime = 0.032 # intTime of -1 = 32 microseconds
				intTime += baseInt #was 550 # compensation for minimum microsecond exposure  0.382, 1200 makes the butterfly scan gradients look most smooth (implying it must be close)
##				le = [0] * pixels
				cieXval = 0.0
				cieYval = 0.0
				cieZval = 0.0
				chlAval = 0.0
				chlBval = 0.0
				nIRval = 0.0
				nUVval = 0.0
				
				#ts = "Radiance W/(sr*sqm*nm)\nInt.: "  + str(intTime)  + "ms Scans: " +  str(nScans)
				pan = int((int(output[0]) - panStart) / pan_Res)
				tilt = int((int(output[1]) - tiltStart) / tilt_Res)
				hspecPan[pan] = int(output[0])
##				hspecTilt[tiltDim-1-tilt] = int(output[1])
				hspecTilt[tilt] = int(output[1])
				loc = 0;
				for i in range(0, pixels, boxcarN): #increment in boxcar size
					xSum = 0
					for k in range(0, boxcarN): #repeat boxcar window
						if(i+k < pixels):
							if radSens[i+k] > 0 and len(output) > loc+5 :
								#print(" " + output[loc+5] + ", " + darkVals[loc+boxedLength*j] + ", " + str(radSens[i+k]))
								#x = float(output[loc+5])-float(darkVals[loc+5+boxedLength*j]) # subtract corresponding dark value
								#x = float(output[loc+5])-250 # subtract corresponding dark value
								x = (float(output[loc+5])-float(darkVals[loc+boxedLength*j])) / boxcarN # subtract corresponding dark value
								if(x > 0):
										#linMultiplier = linCoefs[0]*math.log((x+1)*linCoefs[1])
									x = math.exp(math.log(x)*linCoefs[0] + linCoefs[1])
								if(x < 0):
									x = -1 * math.exp(math.log(-x)*linCoefs[0] + linCoefs[1])
										#linMultiplier = -1* linCoefs[0]*math.log((-x+1)*linCoefs[1])
##								x = (x/linMultiplier) / (radSens[i+k] * intTime * wavelengthBins[i+k])
##								x = (x/linMultiplier) / (radSens[i+k] * intTime)
								x = (x) / (radSens[i+k] * intTime)		
##								le[i] = x
								xSum += x
##								if loc == 0 or loc == 1 or loc ==2:
##								if loc == pixels-1:
##									print("dark:" + darkVals[loc+boxedLength*j] + " light:" + output[loc+5] + " sens:" + str(radSens[i+k]) + " x:"+ str(x))
								x = x * wavelengthBins[i+k]
								cieXval += x * cieXt[i+k]
								cieYval += x * cieYt[i+k]
								cieZval += x * cieZt[i+k]
								chlAval += x * chlAt[i+k]
								chlBval += x * chlBt[i+k]
								nIRval += x * nIRt[i+k]
								nUVval += x * nUVt[i+k]
								
								lum += x * cieYt[i+k]
					#hspec[tilt, pan, loc-1] += xSum/boxcarN # CHECK BOXCAR shouldn't be used here
					hspec[tilt, pan, loc] += xSum
					loc += 1
				lum = lum * 683 #luminance: W/(sr*sqm*nm)
				#ts = "Scanning " +  str(float(pan + (tilt * panDim)) / float(tiltDim * panDim) + "% done")
				#print(str(tiltDim) + ", " + str(panDim))
				ts = str(round(float(pan + (tilt * panDim)) / float(tiltDim * panDim) * 100.0)) + "% done"
				#print(ts)
				imLum[tiltDim-1-tilt, pan] = lum

				# convert to sRGB * set white balance to match computer screen
				imRt = 3.24*cieXval -1.54*cieYval - 0.50*cieZval
				imGt = (-0.97*cieXval + 1.88*cieYval + 0.04*cieZval) * 1.44
				imBt = (0.06*cieXval -0.20*cieYval + 1.06*cieZval) *  1.71

				imR[tiltDim-1-tilt, pan] = imRt
				imG[tiltDim-1-tilt, pan] = imGt
				imB[tiltDim-1-tilt, pan] = imBt

				if(imRt > maxRGB):
					maxRGB = imRt
				if(imGt > maxRGB):
					maxRGB = imGt
				if(imBt > maxRGB):
					maxRGB = imBt

				if int(output[4]) > 0:
					imSatR[tiltDim-1-tilt, pan] = 255
				imSatB[tiltDim-1-tilt, pan] = int(output[4])

				imI[tiltDim-1-tilt, pan] = nIRval
				imGG[tiltDim-1-tilt, pan] = cieYval
				imU[tiltDim-1-tilt, pan] = nUVval

				if(nIRval > maxIGU):
					maxIGU = nIRval
				if(cieYval > maxIGU):
					maxIGU = cieYval
				if(nUVval > maxIGU):
					maxIGU = nUVval

				imChlA[tiltDim-1-tilt, pan] = chlAval / (chlAval+nIRval)
				imChlB[tiltDim-1-tilt, pan] = chlBval / (chlBval+nIRval)

				

				ct = time.time()
				#print("time: " + str(tt-ct))
				if ct > tt:
					tt = ct + 1 # time to next plot in seconds
					statusLabel.config(text=ts)
					plotGraph("")
					#root.after(1, plotGraph(ts))

	if updateDuringDark == 1:
		root.after(1, getSpec)
	else:
		getSpec()
		


def togglePreview():
	global preview
	preview += 1
	if preview > 4:
		preview = 1 #reset set to zero if you want to retain delayed plotting

	if preview == 0:
		btPreview.config(relief="sunken")
		btPreview.config(text="Delay")
	if preview == 1:
		btPreview.config(relief="raised")
		btPreview.config(text="RGB")
	if preview == 2:
		btPreview.config(text="Sat.")
	if preview == 3:
		btPreview.config(text="IGU")
	if preview == 4:
		btPreview.config(text="NDVI")

	plotGraph("")
		
##	if btPreview.config("relief")[-1] == "sunken":
##		btPreview.config(relief="raised")
##		preview = 0
##	else:
##		btPreview.config(relief="sunken")
##		preview = 1


def goTL():
	if(scanningFlag == 0):
		btTL["state"] = "disabled"
		ts = "l" + tiltTop.get()
		ser.write(str.encode( ts ))
		serFlag = 0
		while serFlag == 0: # wait for tilt to get to where it's going
			output = ser.readline()
			if(output.startswith(b't')):
				serFlag = 1
		time.sleep(0.1)
		ts = "p" + panLeft.get()
		ser.write(str.encode( ts ))
		serFlag = 0
		while serFlag == 0: # wait for pan to get to where it's going
			output = ser.readline()
			if(output.startswith(b'p')):
				serFlag = 1
		btTL["state"] = "active"

def goTR():
	if(scanningFlag == 0):
		btTR["state"] = "disabled"
		ts = "l" + tiltTop.get()
		ser.write(str.encode( ts ))
		serFlag = 0
		while serFlag == 0: # wait for tilt to get to where it's going
			output = ser.readline()
			if(output.startswith(b't')):
				serFlag = 1
		time.sleep(0.1)
		ts = "p" + panRight.get()
		ser.write(str.encode( ts ))
		serFlag = 0
		while serFlag == 0: # wait for pan to get to where it's going
			output = ser.readline()
			if(output.startswith(b'p')):
				serFlag = 1
		btTR["state"] = "active"

def goBL():
	if(scanningFlag == 0):
		btBL["state"] = "disabled"
		ts = "l" + tiltBot.get()
		ser.write(str.encode( ts ))
		serFlag = 0
		while serFlag == 0: # wait for tilt to get to where it's going
			output = ser.readline()
			if(output.startswith(b't')):
				serFlag = 1
		time.sleep(0.1)
		ts = "p" + panLeft.get()
		ser.write(str.encode( ts ))
		serFlag = 0
		while serFlag == 0: # wait for pan to get to where it's going
			output = ser.readline()
			if(output.startswith(b'p')):
				serFlag = 1
		btBL["state"] = "active"

def goBR():
	if(scanningFlag == 0):
		btBR["state"] = "disabled"
		ts = "l" + tiltBot.get()
		ser.write(str.encode( ts ))
		serFlag = 0
		while serFlag == 0: # wait for tilt to get to where it's going
			output = ser.readline()
			if(output.startswith(b't')):
				serFlag = 1
		time.sleep(0.1)
		ts = "p" + panRight.get()
		ser.write(str.encode( ts ))
		serFlag = 0
		while serFlag == 0: # wait for pan to get to where it's going
			output = ser.readline()
			if(output.startswith(b'p')):
				serFlag = 1
		btBR["state"] = "active"
	
def goZero():
	if(scanningFlag == 0):
		btZero["state"] = "disabled"
		ser.write(str.encode( "l0" ))
		serFlag = 0
		while serFlag == 0: # wait for tilt to get to where it's going
			output = ser.readline()
			if(output.startswith(b't')):
				serFlag = 1
		ser.write(str.encode( "p0" ))
		serFlag = 0
		while serFlag == 0: # wait for pan to get to where it's going
			output = ser.readline()
			if(output.startswith(b'p')):
				serFlag = 1
		btZero["state"] = "active"

def showRes(a,b,c):
	if(scanningFlag == 0):
		try:			
			pan = math.floor( (int(panRight.get()) - int(panLeft.get())) / int(panRes.get()) ) +1
			tilt = math.floor( (int(tiltTop.get()) - int(tiltBot.get())) / int(tiltRes.get()) ) +1
			pan = math.floor( (int(panTo.get()) - int(panFrom.get())) / int(panResolution.get()) ) +1
			tilt = math.floor( (int(tiltTo.get()) - int(tiltFrom.get())) / int(tiltResolution.get()) ) +1
			ts = str(pan) + "x" + str(tilt)
			statusLabel.config(text=ts)
##			print(ts)
		except:
			return
		


# detect when pan and tilt variables change and show resolution
panTo.trace_add('write', showRes)
panFrom.trace_add('write', showRes)
panResolution.trace_add('write', showRes)
tiltTo.trace_add('write', showRes)
tiltFrom.trace_add('write', showRes)
tiltResolution.trace_add('write', showRes)


def onmouse(event):
	global panDim, tiltDim, hspec, wavelengthBoxcar
	global plotImX, plotImY, selX, selY, refs
	clickX = event.x
	clickY = event.y
##	print("mouse event")
##	print('x:' + str(event.x) + ' y:' + str(event.y))
##	print(  plot_frame.bbox(plot) )
	imAR = panDim/tiltDim # aspect ratio h/w y/x
	frameAR = plotImX/plotImY # width is 2, height is 3
	padding = 2
	if(imAR > frameAR): ## frame narrower than image, image width scales with frame width
		scale = (panDim)/(plotImX)
	else: ## frame wider than image, image height scales with frame height
		scale = (tiltDim)/(plotImY)

	centreX = plotImX/2
	centreY = plotImY/2
	selX = int(math.floor( (panDim/2) + scale*(clickX-centreX)  )) ## offsets between click and centre of image
	selY = int(math.floor( (tiltDim/2) + scale*(clickY-centreY) ))
	selY = tiltDim-selY-1


##	print("Selection:" + str(selX) + ", " + str(selY))
	#---ensure selected coordinates match image dimensions---
	if(selX < 0):
		selX = 0
	if(selX > panDim-1):
		selX = panDim-1
	if(selY < 0):
		selY = 0
	if(selY > tiltDim-1):
		selY = tiltDim-1


	if(len(hspec)>0):
##		print("plot update")
		le = hspec[selY][selX]
		if(reflFlag == 1):
			with np.errstate(invalid='ignore'):
				le = le*100*refs
		[ax[i].clear() for i in range(1)]
		ax[0].plot(wavelengthBoxcar,le)
		ax[0].set_ylim(ymin=0)
		canvas.draw()

##	tR = imR[tiltDim-1-selY, selX]
##	tG = imG[tiltDim-1-selY, selX]
##	tB = imB[tiltDim-1-selY, selX]
##	print("R:"+str(tR)+" G:"+str(tG)+" B:"+str(tB))


def loadFile():
	global fileImportFlag, loadPath, maxRGB, maxIGU
	filetypes = (
		('Hyperspec Files', '*.csv'),
		('All files', '*.*')
	)

	try:
		loadPath = fd.askopenfilename(
			title='Open a file',
			initialdir='./scans/',
			filetypes=filetypes)
		print('Loading: ' + loadPath)
		fileImportFlag = 1
		maxRGB = 1E-10
		maxIGU = 1E-10
		getSpec()
	except:
		fileImportFlag = 0
		return

def setReflVal():
	global reflVal, reflFlag, selX, selY, refs, hspec, imR, imG, imB, maxRGB, wbR, wbG, wbB, wbI, wbGG, wbU, maxIGU, tiltDim
	if(reflFlag == 0):
		reflVal = setRefl.get()
		print("Reflectance standard set to " + reflVal + "%")
##		try:
		if(float(reflVal) > 0 and len(hspec) > 0 and selX != -1 and selY != -1):
			refs = hspec[selY][selX]
##			for i in range(0, len(refs)):
##				if(refs[i] != 0):
##					refs[i] = (float(reflVal)/100)/refs[i]
			with np.errstate(divide='ignore', invalid='ignore'):
				refs = (float(reflVal)/100)/refs # can't deal with zeros
					
##			m = refs
##			m = float(reflVal)/100
##			refs = np.divide(m, refs, out=np.zeros_like(m), where=refs!=0)
					
			btRefl.config(text="Clear Ref.")
			reflFlag = 1
			setRefl["state"] = "disabled"

			# adjust image white-balance
			wbR = imR[tiltDim-1-selY, selX]
			wbG = imG[tiltDim-1-selY, selX]
			wbB = imB[tiltDim-1-selY, selX]
			wbI = imI[tiltDim-1-selY, selX]
			wbGG = imGG[tiltDim-1-selY, selX]
			wbU = imU[tiltDim-1-selY, selX]
##			print("Raw R:"+str(wbR)+" G:"+str(wbG)+" B:"+str(wbB))

			tmaxRGB = wbR
			if(wbG > tmaxRGB):
				tmaxRGB = wbG
			if(wbB > tmaxRGB):
				tmaxRGB = wbB
				
			wbR = tmaxRGB/wbR
			wbG = tmaxRGB/wbG
			wbB = tmaxRGB/wbB

			tmaxIGU = wbI
			if(wbGG > tmaxIGU):
				tmaxIGU = wbGG
			if(wbU > tmaxIGU):
				tmaxIGU = wbU
				
			wbI = tmaxIGU/wbI
			wbGG = tmaxIGU/wbGG
			wbU = tmaxIGU/wbU
##			print("Multiplier R:"+str(wbR)+" G:"+str(wbG)+" B:"+str(wbB))
##				
			plotGraph("Reflectance set")

		else:
			reflFlag = 0
			print("Reflectance cleared")
##		except:
##			reflFlag = 0
##			print("Error reading ref value")

	else:
		clearRefl()
		# clear reflectance


def clearRefl():
	global reflFlag, wbR, wbG, wbB, wbI, wbGG, wbU, hspec, le
	btRefl.config(text="Set Ref.%")
	reflFlag = 0
	setRefl["state"] = "normal"
	wbR = 1.0 # reset white balance
	wbG = 1.0
	wbB = 1.0
	wbI = 1.0
	wbGG = 1.0
	wbU = 1.0
	if(len(hspec)>0):
##		print("plot update")
		le = hspec[selY][selX]
		[ax[i].clear() for i in range(1)]
		ax[0].plot(wavelengthBoxcar,le)
		canvas.draw()
	plotGraph("Reflectance cleared")


def startStop():
	global stopFlag, scanningFlag
	if(scanningFlag == 0 and stopFlag == 0):
		getSpec()
		return
	if(scanningFlag == 1 and stopFlag == 0):
		print("Stopping")
		stopFlag = 1
		getSpec()
		return
	


##updatePlotRes() # run at start to set image size

##------------TOP FRAME-------------
## ┌ ┐└ ┘ ╬ □ ○
	
frame1 = Frame(root)
frame1.grid(row=0, column=0, sticky=N+E+W)
frame1.rowconfigure(0, weight=1)
frame1.rowconfigure(1, weight=1)
frame1.rowconfigure(2, weight=1)

frame1.columnconfigure(0, weight=0)
frame1.columnconfigure(1, weight=1)
frame1.columnconfigure(2, weight=1)
frame1.columnconfigure(3, weight=1)
frame1.columnconfigure(4, weight=1)


btStart = Button(frame1, text='Start', command= lambda: startStop())
btStart.grid(row=0, column=0, padx=2, pady=2, sticky=N+S+E+W)

btTL = Button(frame1, text="┌", relief="raised", command= lambda: goTL())
btTL.grid(row=0, column=1, padx=2, pady=2, sticky=N+S+E+W)

tiltTop = Entry(frame1, textvariable = tiltTo, width =11)
tiltTop.grid(row=0, column=2, padx=2, pady=2, sticky=N+W)

btTR = Button(frame1, text="┐", relief="raised", command= lambda: goTR())
btTR.grid(row=0, column=3, padx=2, pady=2, sticky=N+S+E+W)

resMsg = Label(frame1, text = "Res.")
resMsg.grid(row=0, column=4, padx=2, pady=2, sticky=S+W)

btLoad = Button(frame1, text="Load", relief="raised", command= lambda: loadFile())
btLoad.grid(row=1, column=0, padx=2, pady=2, sticky=N+S+E+W)

panLeft = Entry(frame1, textvariable = panFrom, width =11)
panLeft.grid(row=1, column=1, padx=2, pady=2, sticky=N+W)

btZero = Button(frame1, text="○", relief="raised", command= lambda: goZero())
btZero.grid(row=1, column=2, padx=2, pady=2, sticky=N+S+E+W)

panRight = Entry(frame1, textvariable = panTo, width =11)
panRight.grid(row=1, column=3, padx=2, pady=2, sticky=N+W)

panRes = Entry(frame1, textvariable = panResolution, width =11)
panRes.grid(row=1, column=4, padx=2, pady=2, sticky=N+W)

btPreview = Button(frame1, text="RGB", relief="raised", command= lambda: togglePreview())
btPreview.grid(row=2, column=0, padx=2, pady=2, sticky=N+S+E+W)

##statusLabel = Label(frame1, text = "Status", fg="gray", justify="left")
##statusLabel.grid(row=2, column=0, padx=(5), pady=5, sticky=N+W)

btBL = Button(frame1, text="└", relief="raised", command= lambda: goBL())
btBL.grid(row=2, column=1, padx=2, pady=2, sticky=N+S+E+W)

tiltBot = Entry(frame1, textvariable = tiltFrom, width =11)
tiltBot.grid(row=2, column=2, padx=2, pady=2, sticky=N+W)

btBR = Button(frame1, text="┘", relief="raised", command= lambda: goBR())
btBR.grid(row=2, column=3, padx=2, pady=2, sticky=N+S+E+W)

tiltRes = Entry(frame1, textvariable = tiltResolution, width =11)
tiltRes.grid(row=2, column=4, padx=2, pady=2, sticky=N+W)

##frame1.grid_propagate(False)

##---------------FRAME2-----------------
frame2 = Frame(root)
frame2.grid(row=1, column=0, sticky=N+E+W)
frame2.rowconfigure(0, weight=1)
frame2.rowconfigure(1, weight=1)

frame2.columnconfigure(0, weight=1)
frame2.columnconfigure(1, weight=1)
frame2.columnconfigure(2, weight=1)
frame2.columnconfigure(3, weight=1)

saveMessage = Label(frame2, text = "Label")
saveMessage.grid(row=0, column=0, padx=2, pady=2, sticky=N+W)

maxIntMessage = Label(frame2, text = "MaxInt(ms)") #μ
maxIntMessage.grid(row=0, column=1, padx=2, pady=2, sticky=N+W)

boxcarMessage = Label(frame2, text = "Boxcar")
boxcarMessage.grid(row=0, column=2, padx=2, pady=2, sticky=N+W)

darkRepMessage = Label(frame2, text = "DarkRep(s)")
darkRepMessage.grid(row=0, column=3, padx=2, pady=2, sticky=N+W)

saveInput = Entry(frame2, textvariable = saveLabel, width =11)
saveInput.grid(row=1, column=0, padx=2, pady=2, sticky=N+S+E+W)

maxInt = Entry(frame2, textvariable = maxIntTime, width =11)
maxInt.grid(row=1, column=1, padx=2, pady=2, sticky=N+W)

boxcar = Entry(frame2, textvariable = boxcarVal, width =11)
boxcar.grid(row=1, column=2, padx=2, pady=2, sticky=N+W)

darkRep = Entry(frame2, textvariable = darkRepVal, width =11)
darkRep.grid(row=1, column=3, padx=2, pady=2, sticky=N+W)

##---------------FRAME3-----------------

frame3 = Frame(root)
frame3.grid(row=2, column=0, sticky=N+E+W)

frame3.columnconfigure(0, weight=1)
frame3.rowconfigure(0, weight=1)

brightnessScale = Scale(frame3, from_=1, to=100, orient='horizontal',command= plotGraph)
brightnessScale.set(100)
brightnessScale.grid(row=0, column=0, padx=2, pady=2, sticky=N+W+E)

statusLabel = Label(frame3, text = "Status", fg="gray", justify="left")
statusLabel.grid(row=0, column=0, padx=2, pady=2, sticky=N+W)


##------------IMAGE FRAME-------------

plot_frame = Frame(root)
plot_frame.grid(row=3, column=0, sticky=N+S+E+W)


plot_frame.columnconfigure(0, weight=1)
plot_frame.rowconfigure(0, weight=1)

plotWidth=400
gridIm = Image.open("grid.png")
gridImt = ImageOps.contain(gridIm, (plotWidth,plotWidth), method=0)
gridImResized = ImageTk.PhotoImage(gridImt)
plot = Label(plot_frame, image = gridImResized, fg="gray", justify="left", cursor= "hand2")

plot.grid(row=0, column=0, padx=0, pady=0, sticky=N+W+E+S)
plot.bind('<1>', onmouse) ## mouse click event
plot_frame.grid_propagate(False)


##------------SPEC PLOT FRAME-------------
spec_frame = Frame(root)
spec_frame.grid(row=4, column=0, sticky=N+S+E+W)

figure = plt.Figure(figsize=(6,3), facecolor='#d9d9d9', tight_layout=True) #figsize=(3,1.5)
canvas = FigureCanvasTkAgg(figure, spec_frame)
canvas.get_tk_widget().grid(row=0, column=0, padx=2, pady=2, sticky=N+S+W)
ax = [figure.add_subplot(1, 1, x+1) for x in range(1)]


spec_frame.columnconfigure(0, weight=1)
spec_frame.columnconfigure(1, weight=0)

##------------REFLECTANCE OUTPUT FRAME-------------
refl_frame = Frame(spec_frame)
refl_frame.grid(row=0, column=1, sticky=N+S+E+W)

setRefl = Entry(refl_frame, textvariable = reflVal, width =8)
setRefl.grid(row=0, column=0, padx=2, pady=2, sticky=N+S+E+W)

btRefl = Button(refl_frame, text="Set Ref.%", command= lambda: setReflVal())
btRefl.grid(row=0, column=1, padx=2, pady=2, sticky=N+S+E+W)

refl_frame.columnconfigure(0, weight=1)
refl_frame.columnconfigure(1, weight=1)


root.rowconfigure(0, weight=0)
root.rowconfigure(1, weight=0)
root.rowconfigure(2, weight=0)
root.rowconfigure(3, weight=1)
root.rowconfigure(4, weight=0)
root.columnconfigure(0, weight=1)

root.bind("<Configure>", updatePlotRes) ## resizing the window calls this function

root.mainloop()
```
