## Supplementary figures and images for "Hyperspectral open source imaging system"

### grid.png

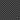

### Print orientations.png

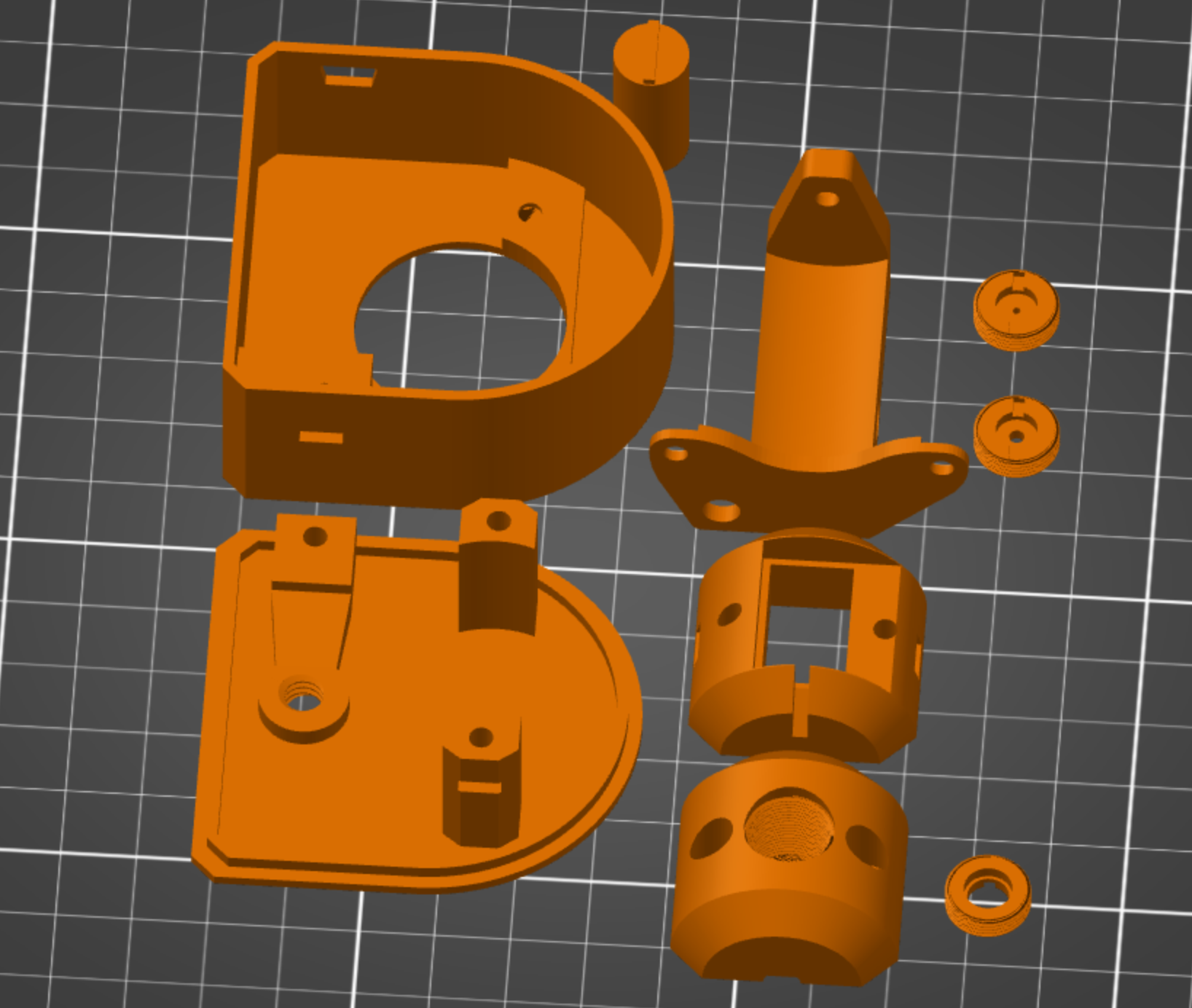
